# Canonical and cross-reactive binding of NK cell inhibitory receptors to HLA-C allotypes is dictated by peptides bound to HLA-C

**DOI:** 10.1101/096958

**Authors:** Malcolm J. W. Sim, Stacy A. Malaker, Ayesha Khan, Janet M. Stowell, Jeffrey Shabanowitz, Mary E. Peterson, Sumati Rajagopalan, Donald F. Hunt, Daniel M. Altmann, Eric O. Long, Rosemary J. Boyton

## Abstract

**Background:** Human natural killer (NK) cell activity is regulated by a family of killer-cell Ig-like receptors (KIR) that bind human leucocyte antigen (HLA) class I. Combinations of KIR and HLA genotypes are associated with disease, including susceptibility to viral infection and disorders of pregnancy. KIR2DL1 binds HLA-C alleles of group C2 (Lys^80^) and KIR2DL2 and KIR2DL3 bind HLA-C alleles of group C1 (Asn^80^). However, this model does not capture allelic diversity in HLA-C or the impact of HLA-bound peptides. The goal of this study was to determine the extent to which the endogenous HLA-C peptide repertoire can influence the specific binding of inhibitory KIR to HLA-C allotypes.

**Results:** The impact of HLA-C bound peptide on inhibitory KIR binding was investigated taking advantage of the fact that HLA-C*05:01 (HLA-C group 2, C2) and HLA-C*08:02 (HLA-C group 1, C1) have identical sequences apart from the key KIR specificity determining epitope at residues 77 and 80. Endogenous peptides were eluted from HLA-C*05:01 and used to test the peptide dependence of KIR2DL1 and KIR2DL2/3 binding to HLA-C*05:01 and HLA-C*08:02 and subsequent impact on NK cell function. Specific binding of KIR2DL1 to the C2 allotype occurred with the majority of peptides tested. In contrast, KIR2DL2/3 binding to the C1 allotype occurred with only a subset of peptides. Cross-reactive binding of KIR2DL2/3 with the C2 allotype was restricted to even fewer peptides. Unexpectedly, two peptides promoted binding of the C2 allotype-specific KIR2DL1 to the C1 allotype. We showed that presentation of endogenous peptides, or predicted HIV Gag peptides, by HLA-C can promote KIR cross-reactive binding.

**Conclusions:** KIR2DL2/3 binding to C1 is more peptide selective than that of KIR2DL1 binding to C2, which provides an explanation for why KIR2DL3–C1 interactions appear weaker than KIR2DL1–C2. In addition, cross-reactive binding of KIR is characterized by even higher peptide selectivity. We demonstrate a hierarchy of functional peptide selectivity of KIR–HLA-C interactions with relevance to NK cell biology and human disease associations. This selective peptide sequence-driven binding of KIR provides a potential mechanism for pathogen as well as self-peptide to modulate NK cell activation through altering levels of inhibition.

## Background

Natural killer (NK) cells are innate lymphocytes with important roles in immune surveillance of cancer and viral infection [1]. NK cell activation is regulated by an array of activating and inhibitory receptors [2]. Inhibitory receptors include CD94-NKG2A, which binds HLA-E, and the killer-cell immunoglobulin-like receptors (KIR) that bind HLA-A, -B and -C molecules [3]. The specificity of inhibitory KIR for different HLA-C allotypes is defined by a dimorphism of a pair of amino acids at position 77 and 80 of HLA-C, whereby KIR2DL1 binds group C2 allotypes (Asn^77^Lys^80^) and KIR2DL2 and KIR2DL3 bind group C1 allotypes (Ser^77^Asn^80^) (Figure 1A). Exchanging amino acids 77 and 80 from a C1 to a C2 allotype, and *vice versa,* revealed that position 80 was the key determinant of KIR specificity for C1 or C2 allotypes [4]. A reciprocal dimorphism exists on the HLA-C specific KIRs at position 44, as KIR2DL2 and KIR2DL3 (Lys^44^) can be converted into C2 binding receptors by mutation to Met^44^, the residue found in KIR2DL1, and *vice versa* [5]. Genetic association studies have highlighted the importance of these interactions, linking combinations of KIR and HLA-C genes in the context of this C1–C2 model (Figure 1A), with multiple disease processes including susceptibility to infectious, autoimmune, and inflammatory disease, cancer, and disorders of pregnancy [3, 6-16]. Examples include protection against chronic hepatitis C virus (HCV) infection in KIR2DL3 and HLA-C1 homozygotes, and increased risk of pre-eclampsia and other pregnancy related disorders when the foetus carries HLA-C2 [9-11]. KIR2DL3 and HLA-C1 is considered a weaker interaction than KIR2DL1 with HLA-C2, allowing disease associations to be explained through the ‘strength of inhibition’ hypothesis, with weaker inhibitory receptor interactions leading to more NK cell activation and clearance of viral infections such as HCV [3, 6, 7, 9, 12-15]. Understanding the basis for specificity in KIR binding to HLA-C allotypes is also relevant to haematopoietic stem cell transplantation for the treatment of certain leukaemias, as the clinical outcome is improved when donor KIR and recipient HLA-C genes are mismatched according to the C1–C2 model for KIR specificity [16-18].

**Figure 1.**
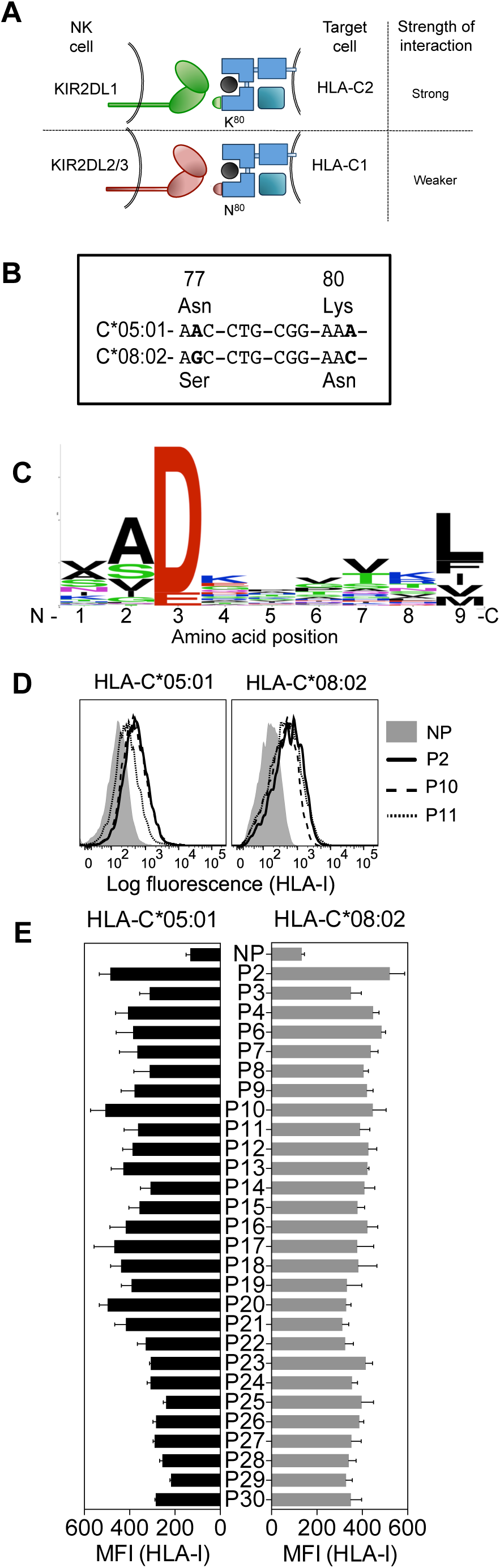
HLA-C*05:01 (group C2) and HLA- C*08:02 (group C1) are almost identical in sequence and HLA-C*05:01-eluted endogenous peptides bind HLA-C*05:01 and HLA-C*08:02. **(A)** Schematic showing how the specificity of inhibitory KIR for different HLA-C allotypes is defined by an amino acid dimorphism at position 77 and 80 of HLA-C, where KIR2DL1 binds group C2 allotypes (Asn^77^Lys^80^) and KIR2DL2 and KIR2DL3 bind group C1 allotypes (Ser^77^Asn^80^) **(B)** Nucleotide sequence alignment of amino acid positions 77-80 of *HLA-C*05:01* and *HLA-C*08:02.* Bold indicates nucleotide differences. **(C)** The motif for HLA-C*05:01 9mer peptides. **(D)** HLA-I expression on 221–C*05:01–ICP47 (left) and RMA-S-C*08:02 *(right)* cells after overnight incubation at 26°C with 100 μM HLA-C*05:01 peptides P2, P10 and P11 and no peptide (NP). **(E)** HLA-I expression on 221–C*05:01–ICP47 (left) and RMA-S-C*08:02 (*right)* cells after overnight incubation at 26°C with 100 mM of 1 of 28 HLA-C*05:01 peptides and no peptide (NP). Mean MFI (median) and SEM of three independent experiments are shown.

KIR binding to HLA-C shows selectivity for peptide, in that certain peptide sequences are incompatible with KIR binding [19-21]. The significance of HLA-C bound peptides in KIR binding for the control of NK cell immune responses has been highlighted by recent examples of HIV escape mutants that result in increased KIR binding due to single amino acid changes in particular viral epitopes [22, 23]. The importance of side chains at positions 7 and 8 of 9 amino acid-long peptides (9mers) was shown using amino acid substituted peptide variants [19, 21, 24, 25]. A study of 60 combinations of amino acid side chains at peptide positions 7 and 8 demonstrated that most combinations reduced KIR2DL2 and KIR2DL3 binding to HLA-C*01:02 (C1) compared to the parent peptide eluted from HLA-C [19]. There is to date no established set of rules to determine the basis for peptide selectivity of KIR for binding HLA-C, although negatively charged residues at positions 7 and 8 are poorly tolerated by KIR [19, 21].

Along with canonical binding to C1, KIR2DL2 and KIR2DL3 display cross-reactive binding to C2 allotypes [26-28]. In contrast, KIR2DL1 is highly specific for C2 allotypes and no functional KIR2DL1 cross-reactivity with C1 allotypes has been reported [27-29]. To determine the extent to which the endogenous HLA-C peptide repertoire can influence the specific binding of inhibitory KIR to HLA-C allotypes, naturally processed peptides were eluted from HLA-C*05:01 and sequenced. HLA-C*05:01, a classical C2 allotype with a Lys at position 80, has shown particularly strong cross-reactivity with KIR2DL2 and KIR2DL3, despite lacking the group C1 Asn at position 80 [26, 27]. Taking advantage of the fact that HLA-C*05:01 (HLA-C group 2, C2) and HLA-C*08:02 (HLA-C group 1, C1) have identical sequences apart from the key KIR specificity determining epitope residues 77-80, we were able to use the same panel of peptides eluted from HLA-C*05:01 to study the impact of HLA-C bound peptide on inhibitory KIR binding in the following interactions: canonical KIR2DL1–C2 and KIR2DL2/3–C1, and cross-reactive KIR2DL2/3–C2 and KIR2DL1–C1.

We show that the majority of peptides tested facilitated the canonical KIR2DL1–HLA-C*05:01 (C2) interaction whereas a more constrained set of peptides allowed the canonical KIR2DL2/3–HLA-C*08:02 (C1) interaction, indicating that the KIR2DL2/3–C1 interaction was more peptide selective than the KIR2DL1–C2 interaction. KIR2DL2 and KIR2DL3 cross-reactive binding to HLA-C*05:01 (C2) was even more peptide selective, while the KIR2DL1–HLA-C*08:02 (C1) interaction was highly peptide selective, occurring with just two peptides. Our results show that peptide sequence plays a significant role in KIR binding and NK cell function, which could be exploited by pathogens, and suggests that a greater dependence on peptide sequence by KIR2DL2/3 may underlie functional differences with KIR2DL1.

## Results

### KIR2DL2 and KIR2DL3 binding to HLA-C*08:02 (C1) is more peptide selective than KIR2DL1 binding to HLA-C*05:01 (C2)

To study the contribution of HLA-C bound peptides to KIR binding and specificity we investigated the impact of KIR binding to HLA-C presenting identical peptides in the context of a C1 and a C2 allotype. HLA-C*05:01 (C2) is almost identical to HLA-C*08:02 (C1), differing only at the positions (77 and 80) that define the C1/C2 allotypes (Figure 1B, Additional file 1A). Peptides naturally bound to HLA-C*05:01 were isolated from 721.221–C*05:01 (221-C*05:01) cells and sequenced by electrospray mass-spectrometry. We identified 72 peptides of which 46 were 9mers (Table 1). The motif for HLA-C*05:01 bound 9mer peptides showed an acidic residue at position 3 (Asp or Glu) and a predominantly hydrophobic anchor at position 9 (Figure 1C). The frequency of amino acids at each position in 9mers showed diversity of residues at most peptide positions, with positions 3, 2 and 9 showing greater preference for particular amino acids (Figure 1C). All peptides contained an acidic residue (Asp or Glu) at position 3. Position 2 showed a preference for small residues (for example Ala). The C-terminal anchor at position 9 was most commonly Leu, with Phe, Ile, Val and Met also present. Our motif for peptides eluted and sequenced from HLA-C*05:01 was highly similar to a motif for HLA-C*05:01 that had been determined using combinatorial peptide libraries and binding data [30]. As expected, a very similar peptide motif had been determined for HLA-C*08:02 [30].

**Table 1.**
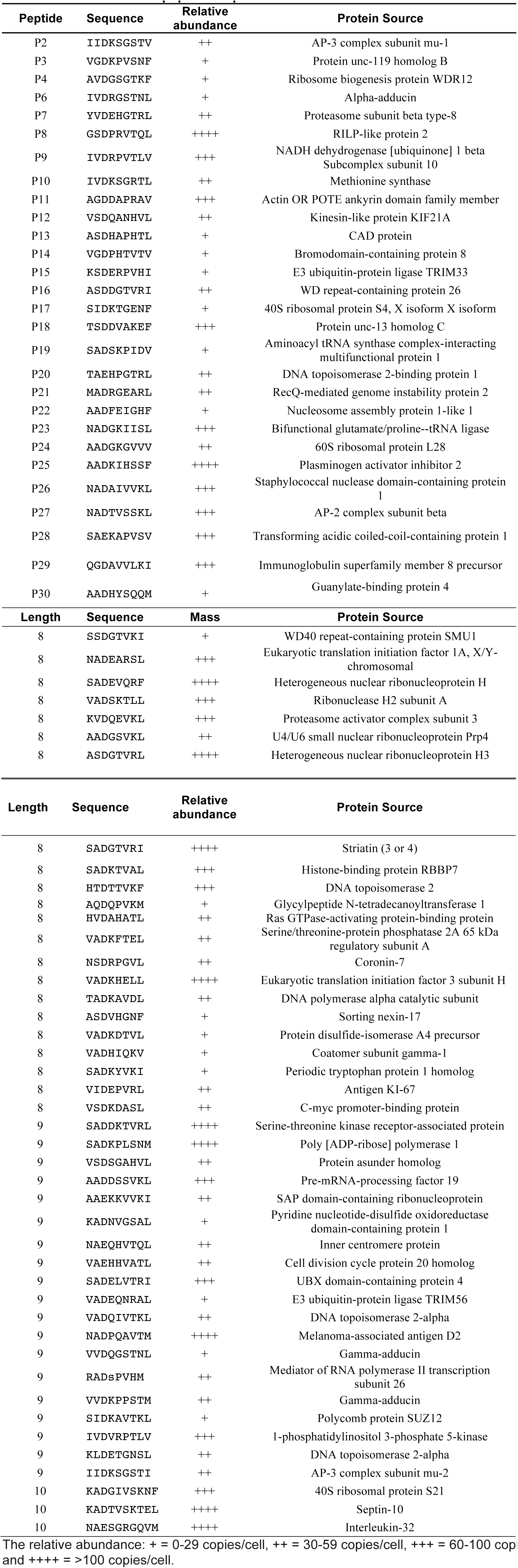
HLA-C*05:01 eluted peptide sequences and relative abundance.

A TAP deficient HLA-C*05:01^+^ cell line was made by expression of the herpes simplex virus-derived TAP inhibitor ICP47 [31] in 221–C*05:01 cells (221–C*05:01-ICP47; Additional file 1B). Of the 46 9mers identified by mass spectrometry, we synthesized the 28 9mer HLA-C*05:01 peptides that included all combinations of amino acids found at positions 7 and 8 (Table 1). The peptide side chains at positions 7 and 8 have the greatest impact on KIR binding [19, 21, 25, 32]. Each of these peptides stabilised HLA-C expression on 221–C*05:01–ICP47 cells after overnight incubation at 26°C, as well as HLA-C*08:02 expression on TAP-deficient RMA-S–C*08:02 cells (Figure 1D, E)[21]. There was a significant positive correlation between stabilization of HLA-C expression on both cell lines and a similar level of expression in the absence of peptide (Figure 1D, E, Additional file 1C). There were a few exceptions, for example peptide 20 (Additional file 1C). However, as most peptides stabilised HLA-C expression on both cell lines to similar extent, we conclude that amino acids 77 and 80 on the otherwise identical HLA-C*05:01 and HLA-C*08:02 do not have a strong influence on peptide loading of TAP-deficient cells. This provided us with the opportunity to compare the binding of KIR2DL1 to a C2 allotype and KIR2DL2 and KIR2DL3 binding to a C1 allotype, using the identical peptide panel.

Soluble KIR-Ig fusion proteins (KIR-Fc) were used to examine KIR2DL1 binding to HLA-C*05:01, and KIR2DL2 and KIR2DL3 binding to HLA-C*08:02 on peptide loaded TAP-deficient cells (Figure 2A,B). Most peptides, 24 of 28, promoted binding of KIR2DL1-Fc to HLA-C*05:01 (C2) (Figure 2A). In contrast, KIR2DL2 and KIR2DL3 binding to HLA-C*08:02 (C1) occurred with only 13 and 11 of 28 peptides, respectively (Figure 2B). This finding remained the same when MFI results were normalised to HLA-I stabilization (KIF-Fc MFI/HLA-I MFI) by each peptide (Additional file 2).

Thus, KIR2DL2 and KIR2DL3 showed greater selectivity for peptides than KIR2DL1 [13/28 and 11/28 compared to 24/28; p<0.002, Chi Square test]. Acidic side chains (Asp or Glu) at position 7 or 8 were present in three of the four peptides that did not promote KIR2DL1 binding, in line with previous studies [19, 21]. These three peptides were also not able to promote KIR2DL2 and KIR2DL3 binding to HLA-C*08:02. Among the peptides that did not confer binding of KIR2DL2 and KIR2DL3, 8 shared a positive charge at position 8 (Lys, Arg, or His), suggesting these amino acids are poorly tolerated by KIR2DL2 and KIR2DL3. Among the 11 peptides that were good for KIR2DL2 and KIR2DL3 binding, seven had Ser or Thr at position 8 and the largest residue found at position 8 was Leu, suggesting a preference for small amino acids. These data showed that KIR2DL2 and KIR2DL3 are more stringent than KIR2DL1 in selection of peptide sequence for binding to HLA-C. The greater selectivity of KIR2DL2 and KIR2DL3 for peptide sequences seems to contradict their reported permissive, cross-reactive binding to group C2 HLA-C allotypes [26-28]. The data suggest a fundamental difference between KIR binding to C1 and C2 allotypes. We next examined the contribution of peptide to KIR2DL2 and KIR2DL3 cross-reactive binding to HLA-C*05:01 (C2).

**Figure 2.**
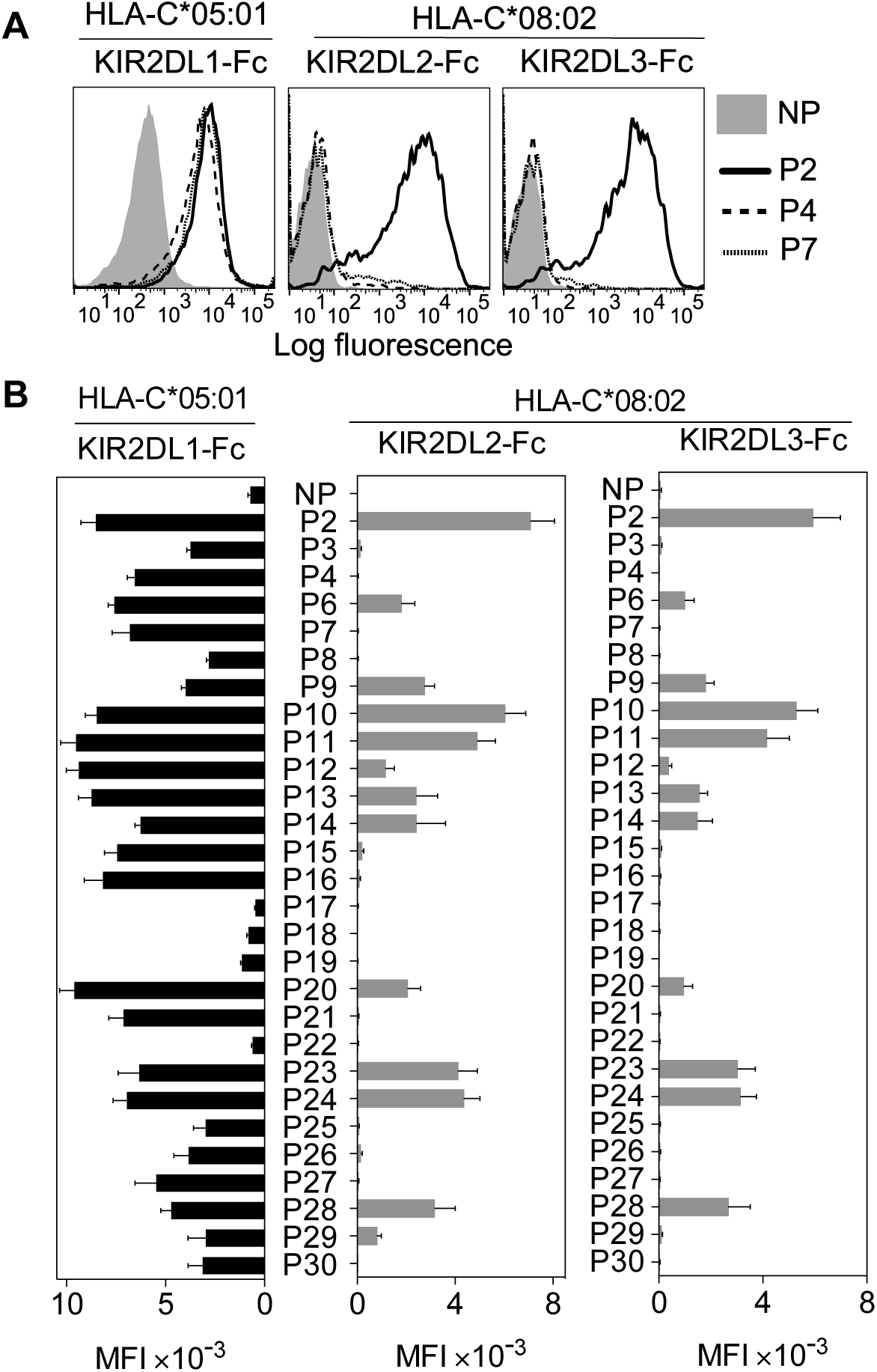
KIR2DL2 and KIR2DL3 binding to HLA-C*08:02 is more peptide selective than KIR2DL1 binding to HLA-C*05:01. **(A)** KIR2DL1-Fc binding to peptide loaded 221–C*05:01–ICP47 *(left),* KIR2DL2-Fc (*middle*) and KIR2DL3-Fc *(right)* binding to RMA-S-C*08:02 cells. Cells were loaded overnight at 26°C with no peptide (NP) or 100 μM HLA-C*05:01 peptides P2, P4 and P7. **(B)** Same as in (A). KIR-Fc binding was assessed after overnight incubation at 26°C with NP or 100 μM of 28 individual HLA-C*05:01 peptides. Mean MFI and SEM of three independent experiments are shown. KIR-Fc (3.6 μg/ml) were conjugated to protein A-alexa 647.

### Cross-reactive KIR2DL2 and KIR2DL3 binding to HLA-C*05:01 (C2) is highly peptide selective

Cross-reactivity of HLA-C*05:01 with KIR2DL2 and KIR2DL3 [27] was confirmed, showing KIR2DL1-Fc, KIR2DL2-Fc and KIR2DL3-Fc binding to 221–C*05:01 cells (Additional file 3A-C). The strength of binding followed the hierarchy KIR2DL1 > KIR2DL2 > KIR2DL3. The same hierarchy was observed with another C2 allotype, HLA-C*04:01 on 221–C*04:01 cells, as reported previously [26, 27], despite weaker overall binding than that observed with 221–C*05:01 cells (Additional file 3A-C). Additionally, an NK cell line expressing KIR2DL3 (YTS–2DL3) was partially inhibited by 221–C*05:01 in cytotoxicity assays, but not by 221–C*04:01 (Additional file 3D,E). Inhibition of KIR2DL3^+^ cells was also tested with primary NK cells, using a gating strategy described in a previous study [33]. Single positive KIR3DL1, single positive KIR2DL1 and single positive KIR2DL3 cells degranulated strongly in response to untransfected 221 cells (Additional file 3F-H). Expression of HLA-C*05:01 on 221 cells had no impact on the degranulation of KIR3DL1-SP cells, while completely inhibiting the degranulation of KIR2DL1-SP NK cells, and partially inhibiting KIR2DL3-SP NK cells (Additional file 3F-H). Thus, HLA-C*05:01 is a functional ligand for KIR2DL1, KIR2DL2 and KIR2DL3.

Next we tested whether KIR2DL2 and KIR2DL3 cross-reactivity resulted from weaker binding across all peptides or greater peptide selectivity. In marked contrast to KIR2DL1-Fc binding to HLA-C*05:01 (Figure 2B), only five peptides conferred cross-reactive binding to KIR2DL2-Fc and KIR2DL3-Fc (Figure 3A,B). Loading of peptides P2, P11, and P24, which have no amino acid in common at positions 7 and 8 (Table 1), resulted in the strongest binding. Thus, KIR2DL2 and KIR2DL3 cross-reactive binding to HLA-C*05:01 was diminished, relative to that of canonical KIR2DL1, due to selective binding to a restricted subset of peptides.

**Figure 3.**
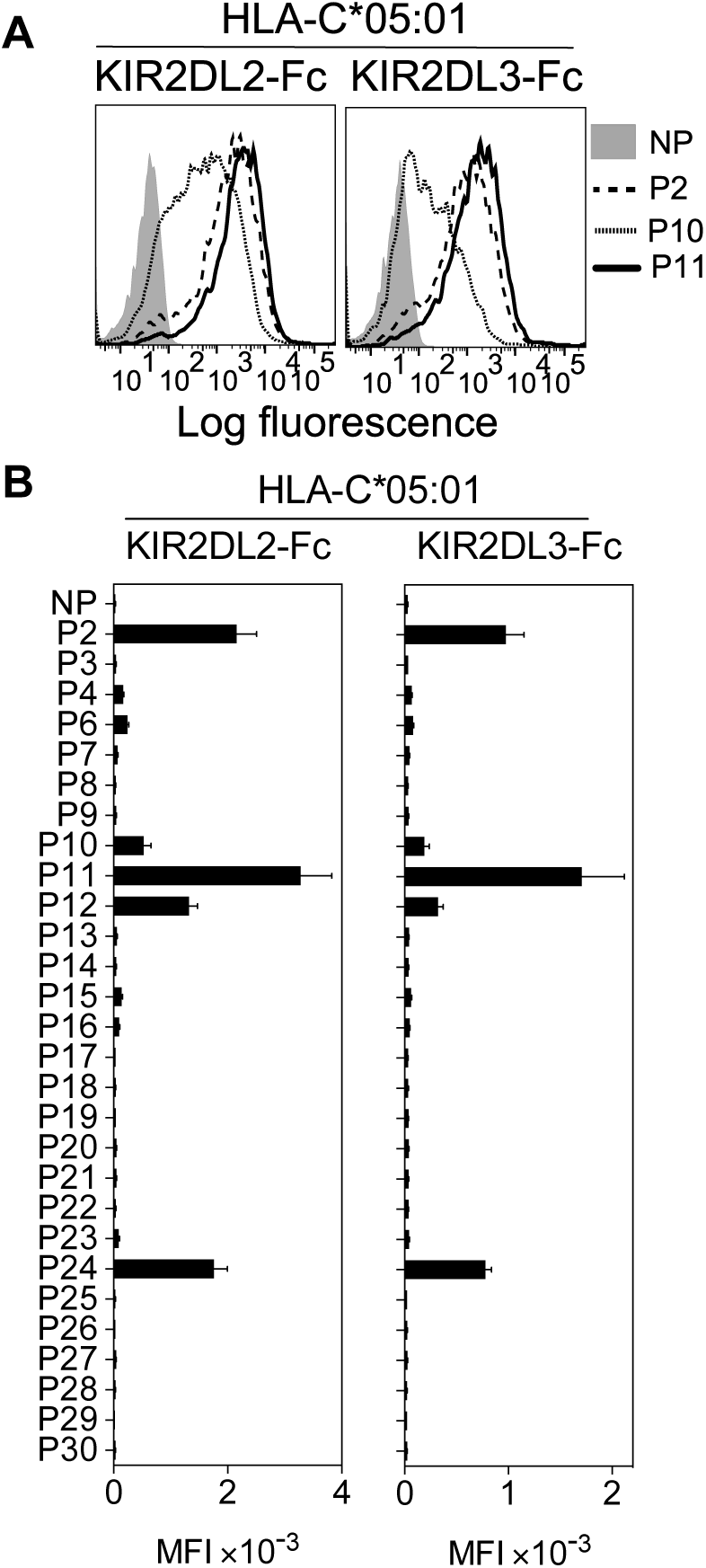
Cross-reactive KIR2DL2 and KIR2DL3 binding to the C2 allotype HLA-C*05:01 is highly peptide selective. **(A)** KIR2DL2-Fc *(left)* and KIR2DL3-Fc *(right)* binding to peptide loaded 221–C*05:01–ICP47, cells. Cells were loaded overnight at 26°C with no peptide (NP) or 100 μM HLA-C*05:01 peptides P2, P10 and P11. **(B)** Same as in (A). KIR-Fc binding was assessed after overnight incubation at 26°C with NP or 100 μM of 28 individual HLA-C*05:01 peptides. Mean MFI and SEM of three independent experiments are shown. KIR-Fc (3.6 μg/ml) were conjugated to protein A-alexa 647.

Peptides that can promote cross-reactive KIR binding should occur by chance for every HLA-C allotype and can originate from endogenous host peptides as well as microbial pathogens. As an example, we tested five peptides derived from HIV Gag that stabilized HLA-I expression on 221–C*05:01–ICP47 cells. Two of them (Gag^60-68^ and Gag^310-318^) promoted canonical binding of KIR2DL1, and one (Gag^310-318^) promoted cross-reactive binding of KIR2DL2 and KIR2DL3 (Additional file 4A-C). These data provide a proof of principle that peptides of viral origin, similar to endogenously processed peptides, can promote cross-reactive KIR binding.

### Peptide sequence determines canonical and cross-reactive KIR2DL2 and KIR2DL3 binding

We then compared the contribution of peptide sequence to canonical and cross-reactive KIR2DL2 and KIR2DL3 binding to HLA-C*08:02 and HLA-C*05:01, respectively (Figure 4). Peptide 2 (P2; IIDKSGSTV) promoted strong KIR2DL2 and KIR2DL3 binding to HLA-C*08:02 and HLA-C*05:01 (Figure 2B, Additional file 2). The role of peptide positions 7 and 8 in KIR binding to P2-loaded HLA-C was explored using amino acid substituted peptides, and binding was compared to that of canonical KIR2DL1 with HLA-C*05:01 (Figure 4). All peptides with amino acid substitutions at positions 7 and 8 had a similar capacity to stabilize HLA-C expression as the parent peptide (Additional file 5A-C). We first tested the impact of decreasing the size of amino acid side chains by substituting Ser at position 7 and Thr at position 8 of P2 with Ala, either singly (IIDKSG**<u>A</u>**TV and IIDKSGS**<u>A</u>**V) or both (IIDKSG**<u>AA</u>**V). Each Ala substituted peptide promoted enhanced and additive cross-reactive binding of KIR2DL2-Fc and KIR2DL3-Fc to HLA-C*05:01, with a greater contribution from an Ala substitution at position 8 than position 7. In contrast, these substitutions had no impact on canonical binding to HLA-C*08:02 (Figure 4A). The strong binding promoted by IIDKSG**<u>AA</u>**V was not due simply to the small side chain size, as substitutions with Gly (IIDKSG**<u>GG</u>**V) abolished cross-reactive binding to HLA-C*05:01 and substantially reduced canonical KIR2DL2 and KIR2DL3 binding to HLA-C*08:02 (Figure 4A). The impact of increased side chain size at position 8 was then tested by replacing position 8 in P2-AA (IIDKSG**<u>AA</u>**V) with Val (P2-AV; IIDKSG**<u>AV</u>**V) and Tyr (P2-AY; IIDKSG**<u>AY</u>**V). Val at position 8 had little impact on KIR2DL2 and KIR2DL3 binding, whereas Tyr at position 8 almost abrogated binding, suggesting that a bulky side chain impeded binding. Consistent with these results, a previous study showed that KIR2DL2 binding to HLA-C*03:04 (C1) was sensitive to side chain size at position 8, with a preference for side chains no larger than Val [25].

**Figure 4.**
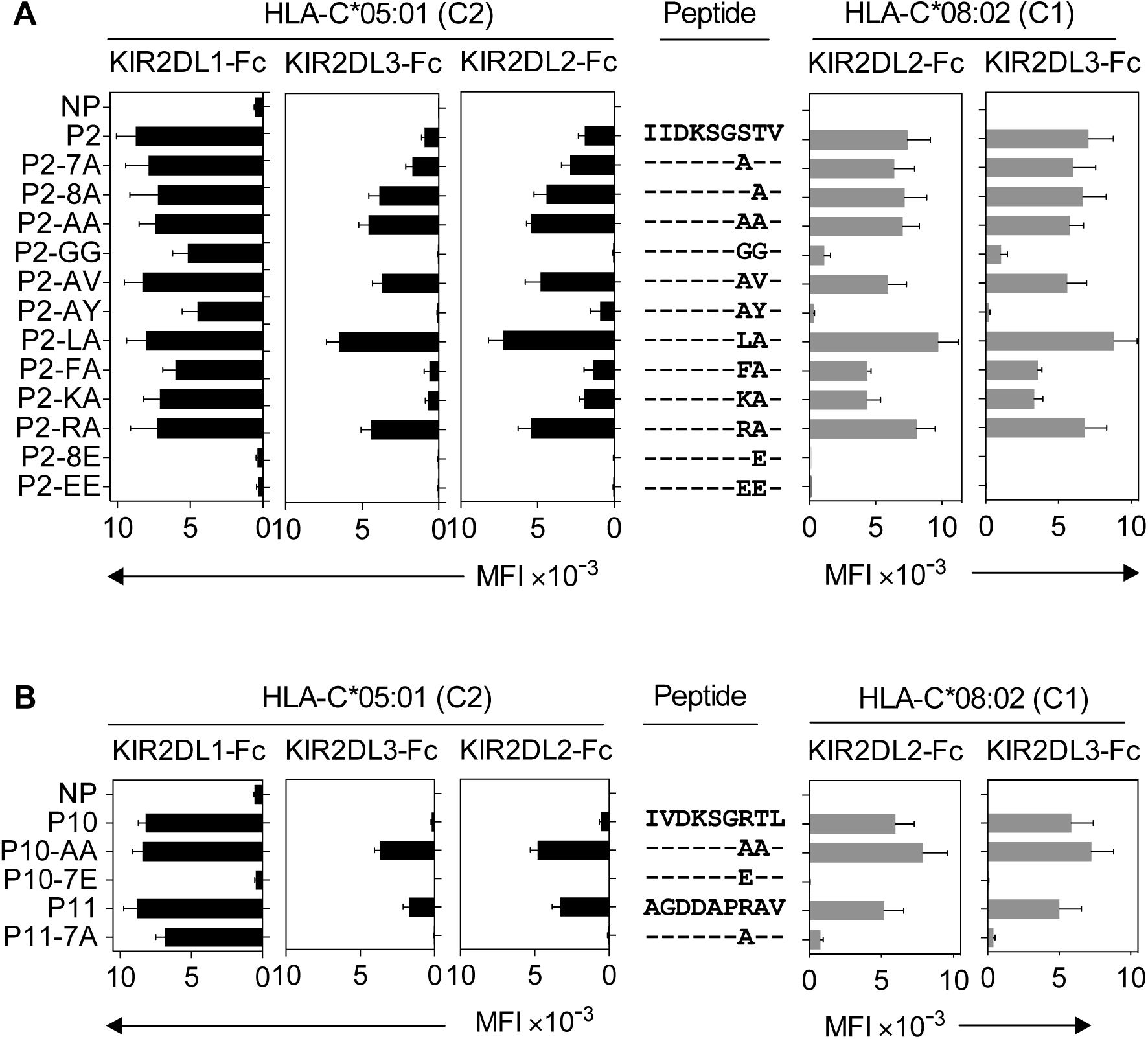
Canonical and cross-reactive KIR2DL2 and KIR2DL3 binding is controlled by amino acids at positions 7 and 8 within the context of a given peptide. **(A)** KIR2DL1-Fc, KIR2DL2-Fc and KIR2DL3-Fc binding to peptide loaded 221–C*05:01–ICP47 (left, black) and KIR2DL2-Fc and KIR2DL3-Fc (right, grey) binding to RMA-S-C*08:02 cells. Cells were loaded overnight at 26°C with no peptide (NP) or 100 μM of peptide P2 (P2-IIDKSGSTV) and amino acid variants of peptide 2 with substitutions at positions 7 and 8. Amino acid sequence substitutions at positions 7 and 8 are shown. Mean MFI and SEM of three independent experiments are shown. KIR-Fc (3.6 μg/ml) were conjugated to protein A-alexa 647. **(B)** KIR2DL1-Fc, KIR2DL2-Fc and KIR2DL3-Fc binding to peptide loaded 221–C*05:01–ICP47 (left, black) and KIR2DL2-Fc and KIR2DL3-Fc (right, grey) binding to RMA-S-C*08:02 cells. Cells were loaded overnight at 26°C with no peptide (NP) or 100 μM of peptides P10 (P10-IVDKSGRTL), P11 (P11-AGDDAPRAV) and amino acid variants of P10 and P11 with substitutions at positions 7 and 8. Amino acid sequence at positions 7 and 8 are shown. Mean MFI and SEM of three independent experiments are shown. KIR-Fc (3.6 μg/ml) were conjugated to protein A-alexa 647.

In a previous study, KIR2DL2 binding to HLA-C*01:02 was strongest with peptides having Leu or Phe at position 7 [19]. We tested KIR2DL2 and KIR2DL3 binding to P2-AA substituted with Leu (P2-LA; IIDKSG**<u>LA</u>**V) or Phe (P2-FA; IIDKSG**<u>FA</u>**V) at position 7 and found that P2-LA promoted the strongest binding of all peptides tested when presented on HLA-C*08:02 or HLA-C*05:01. In contrast, P2-FA promoted weaker binding, a decrease which was more marked in cross-reactive KIR2DL2 and KIR2DL3 binding to HLA-C*05:01, than in canonical binding to HLA-C*08:02.

The strong cross-reactive binding of KIR2DL2 and KIR2DL3 to P11 (AGDDAPRAV, Figure 3) suggested a role for a positively charged residue at position 7 in binding. We tested this by substituting position 7 of P2-AA with Lys (P2-KA; IIDKSG**<u>KA</u>**V) or Arg (P2-RA; IIDKSG**<u>RA</u>**V). While P2-RA promoted strong KIR2DL2 and KIR2DL3 binding, P2-KA was weaker, suggesting that a positive charge alone is not sufficient for strong binding (Figure 4B). Replacement of position 8 with Glu (P2-8E; IIDKSGS**<u>E</u>**V) or positions 7 and 8 with Glu (P2-EE; IIDKSG**<u>EE</u>**V) completely abrogated KIR2DL2 and KIR2DL3 binding to both HLA-C*08:02 and HLA-C*05:01 (Figure 4B). This is consistent with our earlier results (Figure 2) and previous studies demonstrating that negatively charged sides chains at positions 7 or 8 are not competent for binding [19, 21], which is likely due to charge repulsion with the negatively charged KIR binding face [24]. In sharp contrast to KIR2DL2 and KIR2DL3 binding, canonical KIR2DL1 binding to HLA-C*05:01 was largely resistant to changes in amino acid sequence at positions 7 or 8 except to those with acidic residues at positions 7 or 8, similar to our results (Figure 2) and previous studies [21].

### Contribution of peptide sequence at positions 7 and 8 to KIR binding is dependent on peptide context

Canonical and cross-reactive KIR2DL2 and KIR2DL3 binding showed some similarities in preference for amino acids at positions 7 and 8. However, peptide 10 (P10; IVDKSGRTL) promoted strong KIR2DL2 and KIR2DL3 binding to HLA-C*08:02, but weaker cross-reactive binding to HLA-C*05:01 (Figure 2B, Additional file 2). In contrast, peptide 11 (P11; AGDDAPRAV) with similar side chains at positions 7 and 8 conferred canonical and crossreactive binding. P10 and P11 share an Arg at position 7 and have small amino acids at position 8, both of which are compatible with canonical and cross-reactive binding (Figure 4). These results suggest that factors other than side chains at positions 7 and 8 impact on KIR2DL2 and KIR2DL3 binding. To examine this further, residues at positions 7 and 8 of P10 and P11 were substituted with Ala (P10-AA; IVDKSG**<u>AA</u>**L and P11-7A; AGDDAP**<u>A</u>**AV), a combination that promoted strong canonical and cross-reactive binding in the context of P2 (Figure 4A). These substitutions had opposite effects for P10 and P11 (Figure 4B). P10-AA was similar to P2-AA, promoting strong cross-reactive binding and maintaining strong canonical binding. In contrast, P11-7A promoted almost no KIR2DL2 and KIR2DL3 binding, be it canonical or cross-reactive (Figure 4B). In summary, Arg at position 7 is competent for KIR2DL2 and KIR2DL3 crossreactive binding in the context of P2 or P11, but not P10. Ala at position 7 and 8 were competent for KlR2DL2 and KIR2DL3 binding in the context of P2 and P10, but not P11. Therefore, KIR2DL2 and KIR2DL3 binding to HLA-C*05:01 and HLA-C*08:02 appears to be dependent on side chains at position 7 and 8, but the contribution of specific amino acids at position 7 and 8 is valid only in the context of a given 9mer. In contrast, KIR2DL1 binding to HLA-C*05:01 is strong in the presence of the majority of peptide variants accommodating various amino acid side chains, apart from those containing acidic residues (Figure 4 A,B). Overall, the impact of amino acids at position 7 and 8 on KIR2DL2 and KlR2DL3 binding was similar in the context of HLA-C*08:02 and HLA-C*05:01 with a strong correlation in the presence of the same peptides (r = 0.8712, p < 0.0001, Additional figure 6D). Cross-reactive binding of KIR2DL2 and KIR2DL3 was more peptide selective than canonical binding.

Together, these data provide support for an argument that KIR2DL2 and KIR2DL3 canonical binding to HLA-C*08:02 is more peptide selective than that of KIR2DL1 to HLA-C*05:01, that cross-reactive binding to HLA-C*05:01 is even more peptide selective, and that binding is dependent on a combination of different side chains at positions 7 and 8 within a given peptide sequence context.

### High peptide selectivity in functional inhibition of KIR2DL3^+^ NK cells by cross-reactivity with HLA-C*05:01 (C2)

We tested the capacity of KIR2DL2 and KIR2DL3 cross-reactive peptides to inhibit KIR2DL3^+^ NK cells when presented on HLA-C*05:01. YTS cells expressing KIR2DL3 (YTS–2DL3) were tested for inhibition by 221–C*05:01–ICP47 cells that had been loaded with peptides. Inhibition of YTS–2DL3 cytotoxicity correlated well with the binding of KIR2DL3-Fc to HLA-C*05:01 (Figure 5). Strong inhibition was obtained with P2-AA (IIDKSG**<u>AA</u>**V), and partial inhibition with P2 (Figure 5A,B). Loading of peptides that did not promote KIR2DL3 binding to HLA-C*05:01 (P2-GG, IIDKSG**<u>GG</u>**V; P2-EE, IIDKSG**<u>EE</u>**V; or P7, YVDEHGTRL) did not result in inhibition (Figure 5A, B). Inhibition of primary human NK cells was tested in degranulation assays. KIR2DL3-SP NK cells degranulated strongly in response to 221–C*05:01–ICP47 cells that had not been loaded with peptide (Figure 5C, D). 221–C*05:01–ICP47 cells loaded with weaker binders such as P2, P11, or P10 induced partial inhibition (Figure 4, 5D), while those loaded with strong binders such as P2-LA (IIDKSG**<u>LA</u>**V), P2-AA (IIDKSG**<u>AA</u>**V), or P10-AA (IVDKSG**<u>AA</u>**L) induced strong inhibition (Figure 4, 5D). While 221–C*05:01–ICP47 cells loaded peptides that did not promote binding such as P2-GG (IIDKSG**<u>GG</u>**V), P2-EE (IIDKSG**<u>EE</u>**V), P11-7A (AGDDAP**<u>A</u>**AV), and P10-7E (IVDKSG**<u>E</u>**TL) induced as much degranulation as cells in the absence of peptide (Figure 4, 5D). There was a strong inverse correlation between KIR2DL3-Fc binding to, and degranulation of KIR2DL3-SP NK cells in response to 221–C*05:01–ICP47 cells loaded with the same peptides (Figure 5E).

**Figure 5.**
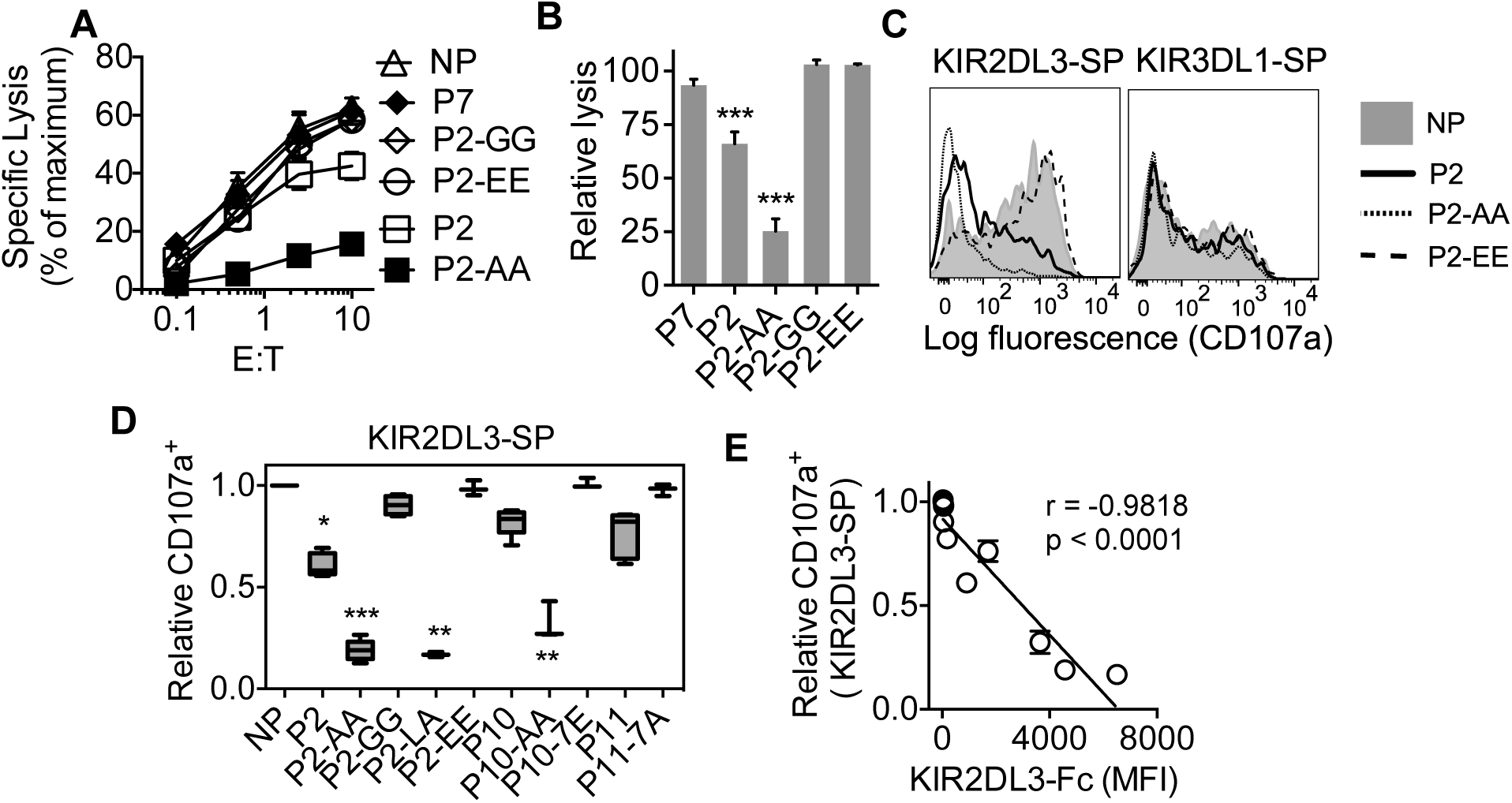
High peptide selectivity, cross reactive binding and functional inhibition of KIR2DL3^+^ NK cells by HLA-C*05:01 (C2). **(A)** Specific lysis of 221–C*05:01–ICP47 cells at different E:T by YTS-2DL3 NK cells. 221–C*05:01–ICP47 target cells were used after overnight culture 26°C in the presence of no peptide (NP), P2, P2-AA, P2-EE, P2-GG, and P7. **(B)** Relative specific lysis of 221–C*05:01–ICP47 cells by YTS-2DL3 NK cells compared to NP at 10:1 (E:T). Mean and SEM of three independent experiments are shown. ***=p<0.001 by students t test. **(C)** Flow cytometry histograms showing degranulation (CD107a expression) of KIR2DL3-SP *(left)* and KIR3DL1-SP *(right)* NK cells in response to 221–C*05:01–ICP47 cells loaded with P2, P2-AA, P2- EE or NP. **(D)** Box plots showing relative expression of CD107a on KIR2DL3-SP NK cells in response to 221–C*05:01–ICP47 cells loaded with P2, P2-AA, P2-EE, P2-GG, P2-LA, P11, P11-7A, P10, P10-AA, P10-7E or NP. N = 3-5. CD107a expression was normalized to value obtained in response to NP for each donor. ***=p<0.001, **=p<0.01, *=p<0.05 by Kruskal-Wallis test with Dunn’s multiple comparisons test. **(E)** KIR2DL3-Fc binding to 221–C*05:01–ICP47 cells correlated with degranulation (CD107a+) of KIR2DL3-SP NK cells in response to 221–C*05:01–ICP47 cells in the presence of P2, P2-AA, P2-EE, P2-GG, P2-LA, P11, P11-7A, P10, P10-AA, P10-7E or NP. Spearman correlation r = −0.955, p<0.0001. CD107a expression was normalized to value obtained in response to NP for each donor.

As an internal control, degranulation of KIR3DL1-SP NK cells was not impacted by the presence of any of the peptides (Figure 5C and Additional file 6). Thus, we have shown that the group C2 allotype, HLA-C*05:01 loaded with peptide that confers binding to KIR2DL3 induces functional inhibition of KIR2DL3+ NK cells.

### Functional cross-reactive KIR2DL1 binding to the C1 allotype, HLA-C*08:02

KIR2DL1 is highly specific for the C2 allotypes [3]. We put this specificity to a test by examining KIR2DL1 binding to HLA-C*08:02 (C1) after loading with the same peptide panel that had been used to test canonical binding to HLA-C*05:01. We found two peptides that promoted KIR2DL1 binding to HLA-C*08:02 (C1), P10 (IVDKSGRTL) and P11 (AGDDAPRAV) (Figure 6A). For P10, this KIR2DL1 cross-reactive binding was about 2- to 3-fold weaker than KIR2DL3 binding to HLA-C*08:02 (Figure 2 and 6A). P10 and P11 share an Arg at position 7 and KIR2DL1 binding to HLA-C*08:02 was lost when Arg at position 7 of P11 was replaced by Ala (P11-7A; AGDDAP**<u>A</u>**AV) (Figure 6A, B). To further test whether Arg at position 7 contributed to crossreactive binding of KIR2DL1 to HLA-C*08:02 an Arg was substituted for the Ala at position 7 of the non-binding peptide P2-AA (IIDKSG**<u>AA</u>**V). The resulting P2-RA (IIDKSG**<u>RA</u>**V) peptide promoted binding of KIR2DL1 to HLA-C*08:02 (Figure 6A, B). This Arg-dependent gain-of-function was not reproduced with a Lys at position 7 (P2-KA; IIDKSG**<u>KA</u>**V, in Figure 6A, B), suggesting specific recognition of Arg at position 7 by KIR2DL1 in the context of HLA-C*08:02.

**Figure 6.**
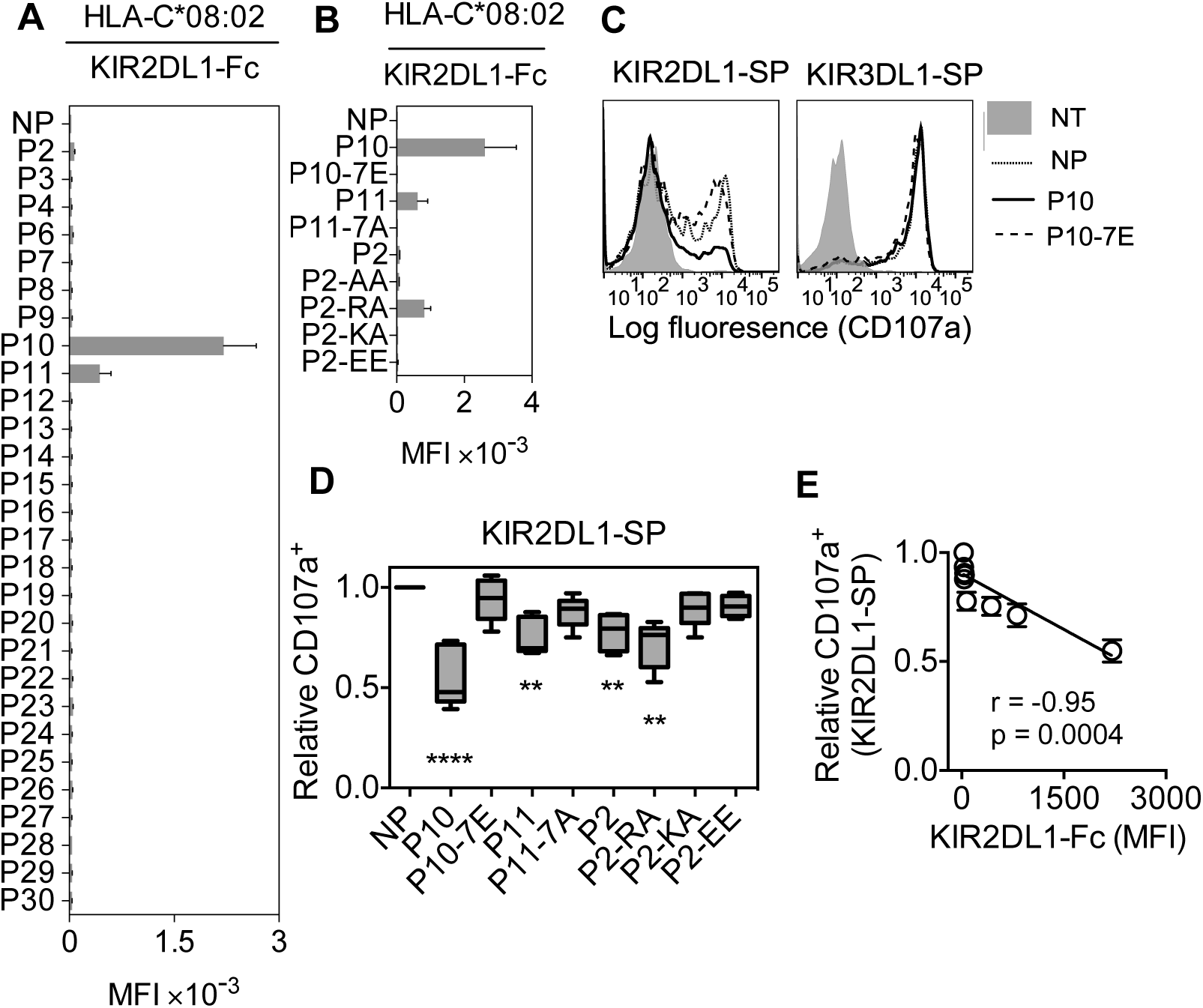
High peptide selectivity and functional cross-reactive binding of KIR2DL1 to the C1 allotype HLA-C*08:02. **(A)** KIR2DL1-Fc binding to RMA-S-C*08:02 cells after overnight incubation at 26°C with NP or 100 μM of 28 individual HLA-C*05:01 peptides. Mean MFI and SEM of three independent experiments are shown. **(B)** KIR2DL1-Fc binding to RMA-S-C*08:02 cells after overnight incubation at 26°C with NP or 100 μM of peptides P10, P10-7E, P11, P11-7A, P2, P2-AA, P2-RA, P2-KA and P2-EE. Mean MFI and SEM of three independent experiments are shown. KIR-Fc (3.6 mg/ml) were conjugated to protein A-alexa 647. **(C)** Flow cytometry histograms showing degranulation (CD107a expression) of KIR2DL1-SP *(left)* and KIR3DL1-SP *(right)* NK cells in response to 221–C*08:02–ICP47 cells pre-incubated overnight with P10, P10-7E or no peptide (NP). Expression of CD107a on KIR2DL1-SP NK cells with no target (NT) cells is also shown. **(D)** Box plots showing CD107a expression on KIR2DL1-SP NK cells in response to 221–C*08:02–ICP47 cells pre-incubated overnight with P10, P10-7E, P11, P11-7A, P2, P2-RA, P2-KA, P2-EE or NP. N = 5-8. CD107a^+^ expression is displayed relative to values obtained with NP. ****=p<0.0001, **=p<0.01 by Kruskal-Wallis test with Dunn's multiple comparisons test. **(E)** KIR2DL1-Fc binding to RMA-S-C*08:02 cells is correlated with degranulation (CD107a^+^) of KIR2DL1-SP NK cells in response to 221–C*08:02–ICP47 cells loaded with P10, P10-7E, P11, P11-7A, P2, P2-RA, P2-KA, P2-EE or NP. CD107a expression was normalized to value obtained in response to NP for each donor. Spearman correlation, p=0.0004, r = −0.95.

To test whether cross-reactive KIR2DL1 binding to HLA-C*08:02 resulted in functional inhibition of NK cytotoxicity, we generated 721.221 cells expressing HLA-C*08:02 (221–C*08:02; Additional file 7A, B), which were further transfected with the TAP inhibitor ICP47 (221–C*08:02–ICP47). KIR2DL1-Fc bound 221–C*08:02–ICP47 cells that had been loaded with P10 (IVDKSGRTL) (Additional file 7C). In assays using primary NK cells, KIR2DL1-SP NK cells degranulated in response to 221–C*08:02–ICP47 cells not loaded with peptide (Figure 6C, D). Loading of 221–C*08:02–ICP47 cells with peptide P10 resulted in inhibition of degranulation by KIR2DL1-SP cells, but not KIR3DL1-SP cells (Figure 6C). As expected, replacement of Arg at position 7 with Glu, P10-7E (IVDKSG**<u>E</u>**TL) that impeded binding also eliminated inhibition (Figure 6B-D). We conclude that cross-reactive binding of KIR2DL1 to HLA-C*08:02 loaded with P10 (IVDKSGRTL) resulted in functional inhibition of NK cell degranulation. While P10 was compatible with canonical binding of KIR2DL1, KIR2DL2 and KIR2DL3 (Figure 2) and with cross-reactive KIR2DL1 binding to HLA-C*08:02 (Figure 6A), it formed a less effective ligand for cross-reactive binding of KIR2DL2 and KIR2DL3 to HLA-C*05:01 (Figure 3). Weak inhibition of degranulation by KIR2DL1-SP NK cells was detected with 221–C*08:02–ICP47 cells loaded with peptides that promoted weak KIR2DL1 binding, P11 (AGDDAPRAV) and P2-RA (IIDKSG**<u>RA</u>**V) (Figure 6D). Replacement of the Arg at position 7 of P2-RA by Lys P2-KA (IIDKSG**<u>KA</u>**V) eliminated inhibition, as expected from the loss of binding (Figure 6B). Similar to KIR2DL3 with HLA-C*05:01, there was a strong inverse correlation between KIR2DL1-Fc binding to, and degranulation of KIR2DL1-SP cells in response to 221–C*08:02–ICP47 cells (Figure 6E). Degranulation of KIR3DL1-SP NK cells was not inhibited by any of the peptides (Figure 6C and Additional file 7D). The ratio of KIR-Fc binding (as mean fluorescence intensity) to the extent of degranulation (as a fraction of each SP cell subset) was similar for KIR2DL1 with HLA-C*08:02 and KIR2DL3 with HLA-C*05:01 (Additional file 7E). We conclude that the highly peptide-selective binding of KIR2DL1 to the group C1 allotype HLA-C*08:02 was competent to induce functional inhibition of KIR2DL1^+^ NK cells.

## Discussion

Disease associations with different combinations of KIR and HLA-C genes have been interpreted through the ‘strength of inhibition’ hypothesis, which posits that KIR2DL1 binding to a C2 allotype is a stronger interaction than KIR2DL3 with C1 [3, 6, 8-15, 34] This hypothesis is based on the weaker binding of soluble KIR2DL3-Fc fusion protein to C1 compared to KIR2DL1-Fc binding to C2, using HLA-C^+^ cells or HLA-C molecules attached to beads [27, 28, 35]. Weaker interactions have been interpreted as potentially beneficial in viral infections such as HCV due to reduced inhibition of NK cells in individuals homozygous for C1 and for KIR2DL3 [9]. Conversely, disorders of pregnancy in mothers carrying a C2/C2 fetus and expressing KIR2DL1 on uterine NK cells have been interpreted as being due to stronger inhibition by KIR2DL1 resulting in decreased activation of specialized uterine NK cells that promote vascular remodeling to increase blood supply to the fetus [10, 11, 36, 37]. However, the strength of inhibition hypothesis has to be reconciled with direct measurements of KIR binding to HLA-C in solution. Dissociation constants for KIR2DL1 with C2, and for KIR2DL2 and KIR2DL3 with C1, have been similar, ranging from 6.9 to 11.2 mM [25, 38, 39]. These measurements, obtained with recombinant HLA-C molecules that had been refolded with one peptide at a time, do not capture the complexity of KIR binding to cells expressing HLA-C molecules associated with a large repertoire of peptides. By examining how HLA-C-bound peptides regulate binding of KIR we provide an explanation for the difference between solution binding measurements and strength of inhibition, and demonstrate a hierarchy in the contribution of both HLA-C allotype and peptide sequence to KIR–HLA-C interactions (Figure 7). Our findings have direct implications for NK biology and human disease associations.

**Figure 7.**
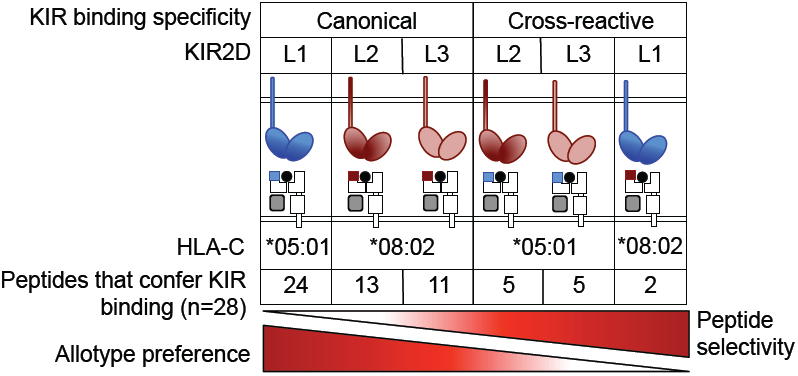
Differences in allotype and peptide selectivity of KIR binding to HLA-C. KIR2DL1, with strong specificity for C2 allotype, exhibits low peptide selectivity in binding C2. KIR2DL2 and KIR2DL3 are more peptide selective than KIR2DL1, and less stringent about allotype preference. KIR2DL2 and KIR2DL3 cross-reactivity with C2 is associated with an even greater peptide selectivity. C2 specific KIR2DL1 cross-reacted with C1 in the context of only 2 peptides. This hierarchy in peptide selectivity of KIR–HLA-C interactions is relevant to NK biology and understanding of disease associations. Peptide sequence-driven binding of KIR to HLA-C provides a potential mechanism for pathogen as well as self-peptide to modulate NK cell activation through altering levels of KIR binding and KIR mediated inhibition.

We took advantage of the sequence identity of HLA-C*05:01 (group C2) and HLA-C*08:02 (group C1), but for the two amino acids that define group C1 and C2 allotypes and specificity of KIR2DL1–C2 and KIR2DL2/3–C1 binding, which allowed us to compare binding of KIR to C1 and C2 allotypes loaded with the same set of peptides. We demonstrated that KIR2DL2/3 binding to C1 is more selective for specific HLA-C–peptide complexes than KIR2DL1–C2, providing an explanation for why KIR2DL3–C1 interactions appear to be weaker than KIR2DL1–C2. KIR2DL1 bound to C2 allotype HLA-C*05:01 in the context of most peptides tested. In addition, we show that the peptide selectivity of KIR2DL2/3 binding was even greater in the context of cross-reactive binding to C2 allotype HLA-C*05:01. Cross-reactive binding of KIR to non-canonical HLA-C allotypes has been described for KIR2DL2 and KIR2DL3 [26-28]. In the crystal structure of a KIR2DL2–HLA-C*03:04 complex the peptide side chains at position 7 and 8 are both orientated upwards towards the KIR and are both making contacts [25], whereas peptide made no direct contribution to binding in the crystal structure of a KIR2DL1–HLA-C*04:01 complex [24]. This is consistent with our data demonstrating a greater contribution of peptide sequence to canonical KIR2DL2/3 binding with C1 than to KIR2DL1 binding to C2.

The peptide sequence-dependent binding of KIR to HLA-C described here contrasts with the prevailing view that peptide selectivity of KIR binding to HLA-C is due mainly to peptide side chains at positions 7 and 8 of 9mers that interfere with binding through their bulk or charge[21, 25]. Furthermore, KIR2DL2 and KIR2DL3 binding is not governed simply by amino acids at positions 7 and 8. For example, replacing Arg-Ala at those positions in the peptide P2 with Ala-Ala did not affect KIR2DL2 and KIR2DL3 canonical binding to C1 or cross-reactive binding to C2. The same Arg-Ala to Ala-Ala substitution in peptide P11 eliminated binding of KIR2DL2 and KIR2DL3 to both C1 and C2 allotype. Finally, peptide P10 with Arg-Thr at positions 7 and 8, which promoted strong KIR2DL2 and KIR2DL3 canonical binding but weak cross-reactive binding, *gained* strong cross-reactive binding to HLA-C*05:01 when changed to Ala-Ala. Thus the contribution of positions 7 and 8 to binding of KIR2DL2 and KIR2DL3 is clearly tied to additional features of the peptide.

KIR2DL1 has strong selectivity for C2 allotypes. Weak cross-reactive binding of KIR2DL1 was reported with group C1 HLA-Cw7 loaded with a single peptide, but was not tested functionally [38, 39]. We show here that two peptides loaded on the C1 allotype HLA-C*08:02 promoted KIR2DL1 binding, which resulted in functional inhibition of KIR2DL1^+^ NK cells. The crystal structure of a canonical KIR2DL1–HLA-C*04:01 complex revealed a binding site made largely of shape complementarity and of electrostatic forces between a positively charged HLA-C molecule and a negatively charged KIR[24]. Lys^80^ of HLA-C*04:01 is accommodated by a specific pocket in KIR2DL1, in which Met^44^, Ser^184^, and Glu^187^ interact directly with HLA-C. The peptide made no direct contribution to binding, which may explain why KIR2DL1 binds to HLA-C*04:01 and, as shown here, to HLA-C*05:01 in the context of most peptides [21, 24]. It is also consistent with the notion that KIR2DL1 and C2 allotypes have co-evolved more recently than KIR2DL2/3 with C1 allotypes as a more stringent KIR–HLA-C combination [29]. Cross-reactive KIR2DL1 binding to HLA-C*08:02 occurred only with peptides carrying Arg at position 7, suggesting that p7 Arg may compensate for the absence of the C2-defining Lys^80^.

Our data suggests a hierarchy in the contribution of both HLA-C allotype and peptide sequence in KIR binding (Figure 7). KIR2DL1, with strong specificity for C2 allotypes, binds C2 in the presence of most peptides. That peptide sequence contributes minimally to KIR2DL1 binding to C2 [21] is consistent with a lack of peptide contacts in the KIR2DL1–HLA-C*04:01 crystal structure[24]. Together with the greater propensity of KIR2DL2/3 to cross-react with C2 than KIR2DL1 with C1, the data suggests a more fundamental difference between KIR2DL2/3 and KIR2DL1 binding to HLA-C, in which selectivity for HLA-C allotype is inversely correlated with selectivity for peptides (Figure 7).

The use of HLA-C*05:01 and HLA-C*08:02 allotypes in our study has made it possible to examine and compare binding of KIR to a C2 and a C1 allotype in the context of the same peptides. As there is a very large number of KIR combinations with HLA-C allotypes due to extensive polymorphism of KIR [29, 40-42] and of HLA-C, canonical and cross-reactive binding of KIR to HLA-C, and the contribution of peptide to binding, may vary among different KIR–HLA-C combinations. KIR2DL2 and KIR2DL3 cross-reactive binding with C2 indeed varies among C2 allotypes [27].

The high polymorphism of the HLA-C peptide binding site is such that different HLA-C allotypes within group C1 or C2 present very different peptide repertoires. Therefore, a specific and conserved recognition of an HLA-C–peptide complex cannot be achieved by innate receptors such as KIR, which bind to a large number of HLA-C allotypes. Peptides that promote cross-reactive KIR binding should occur with a given probability for each HLA-C allotype and originate from endogenous host proteins as well as microbial pathogens. As an example, of five peptides derived from HIV Gag that were predicted to bind HLA-C*05:01, two of them promoted canonical binding of KIR2DL1, one of which promoted cross-reactive binding of KIR2DL2 and KIR2DL3. Similarly, a screen of peptides from two HCV proteins identified a 9mer sequence that promoted stronger binding of KIR2DL3 to HLA-C*03:04 than binding obtained with an endogenous peptide known to be associated with HLA-C*03:04 [43]. These results illustrate the broader principle of immune subversion by intracellular pathogens capable of generating peptides that bind HLA-C and modulate inhibitory KIR binding. Previous studies have reported HIV escape mutations that were associated with the presence of specific KIR genes, suggesting NK-mediated immune pressure on HIV [23]. There was increased KIR binding to, and NK cell inhibition by, cells infected with viruses carrying the KIR-associated escape mutant. One of the HIV escape mutant peptides has been identified as a p24 Gag epitope, which promoted greater KIR2DL3 canonical binding to HLA-C*03:04 than that of the original HIV Gag sequence [22]. It is remarkable that the difference in binding to a single peptide can have an impact on KIR binding to cells that express a large repertoire of HLA-C–peptide complexes. Analysis of peptide repertoires on virus-infected cells are needed to better understand viral evasion through inhibition of NK cells. As we have shown here, the specificity of KIR for HLA-C is such that a single amino acid change in the peptide can be sufficient to cause large differences in KIR binding and KIR-mediated inhibition, including cross-reactive KIR binding to a non-canonical HLA-C allotype.

A peptide that promotes strong KIR cross-reactivity could push NK cells that are normally unlicensed to a more licensed state through non-canonical KIR–HLA-C binding. This could occur in C1/C1 and C2/C2 homozygotes, as in either case single-positive KIR2DL1^+^ NK cells and single-positive KIR2DL2^+^ or KIR2DL3^+^ NK cells, respectively, lack a canonical HLA-C ligand. Immune pressure by NK cells may select for viral peptides that promote cross-reactivity, given that unlicensed NK cells, which include cells with orphan inhibitory receptors, have been shown to provide stronger anti-viral responses than licensed NK cells in MCMV-infected and influenza virus-infected mice, and in EBV-infected humanized mice [44-47]. According to this model, it would be more difficult for pathogens to evade single-positive KIR2DL1^+^ NK cells in C1/C1 individuals than single-positive KIR2DL3^+^ NK cells in C2/C2, as the KIR2DL1–C1 cross-reactive combination is less prone to peptide sequence-dependent modulation. The new insights gained by our study suggest that interpretation of disease associations with KIR and C1 or C2 genes requires a better appreciation of the contribution of HLA-C associated peptides to KIR binding and NK cell function.

## Materials and Methods

### Study Design

The goal of the study was to investigate the contribution of HLA-C bound peptides to canonical and cross-reactive KIR binding to C1 and C2 allotypes. *In vitro* experiments were carried out to test the level of soluble KIR-Fc binding to peptide-loaded HLA-C on TAP-deficient cells by flow cytometry. Experiments were repeated at least three times and KIR-Fc binding was validated using functional experiments carried out with primary human NK cells from at least three healthy volunteers.

### HLA-C peptide sequencing

HLA-C bound peptides were sequenced as described [48, 49]. HLA-C was affinity purified from cell lysates of 1 × 10^10^ 221–C*05:01-His_10_ using a Ni^2+^ column followed by W6/32 column. NHS:N-hydroxysuccinimide-activated Sepharose 4 fast flow (Pharmacia Biotech) was incubated with W6/32 mAb (5 mg protein/mL Sepharose) overnight at room temperature. The column was blocked with 100 mM ethanolamine, pH 9.0, overnight before extensive washing with PBS. Filtered lysates of 221–C*05:01 cells were passed over a pre-column (IgG2a) followed by the W6/32 column. Columns were washed with 20 column volumes of lysis buffer, low salt buffer, high salt buffer and no salt buffer. Lysis buffer was 20 mM Tris HCl pH 8.0 with 150 mM NaCl with 1% CHAPS, 1 mM EDTA, 0.2% sodium azide and protease inhibitors. Low salt buffer was 50 mM Tris HCl with 250 mM NaCl (pH 8.0). High salt buffer was 50 mM Tris HCl with 1 mM NaCl (pH 8.0). No salt buffer was 50 mM Tris-HCl (pH 8.0). HLA-C was eluted from the W6/32 column with 4 column volumes of 0.2 M acetic acid. Peptides were separated by ultrafiltration to remove HLA-C heavy chain and b_2_M (3 kDa, Amicon Ultra-15, Millipore, USA). The filtrate was made up in 10% acetic acid and frozen. 1ml was dried down in a speed vac and reconstituted at 1 × 10^7^ cell equivalents/ml in 0.1% acetic acid. Approximately 2x10^7^ cell equivalents were made up to 4 ml in 0.1% acetic acid and spiked with 100 fmol internal standards angiotensin and vasoactive peptide. Sample was loaded onto an irregular C18 (5-20 mM diameter) capillary pre-column (360 mM outer diameter, 75 mM inner diameter) and washed with 0.1% acetic acid before connecting the pre-column to a C18 analytical capillary column (360 mM outer diameter, 50 mM inner diameter) equipped with an electrospray emitter tip. Peptides were gradient eluted by increasing organic solvent from 100% solvent A with 0% solvent B to 60% solvent B over 60 min (where Solvent A is 0.1 M acetic acid in H_2_O, and solvent B is 70% acetonitrile and 0.1 M acetic acid). Ions were electrosprayed into a highresolution mass spectrometer on elution (Thermo Orbitrap XL, USA). The five most abundant precursor masses present in MS1 were selectively fragmented in a linear ion trap by collision activated dissociation followed by electron transfer dissociation. On completion, the open mass spectrometry search algorithm was used to search the human RefSeq NCBI Reference Sequence Database (http://www.ncbi.nlm.nih.gov/refseq/). Peptides were manually validated.

### Peptide HLA-C stabilization assays

Peptide stabilization of HLA-C was assessed by flow cytometry largely as described [19, 21].1.25 × 10^5^ cells were incubated overnight at 26°C with 100 μM of synthetic peptide, as indicated. The following day cells stained with PE-Cy5 HLA-A,B,C mAb (G46-2.6, BD) at 4°C. Peptides were synthesized GL Biochem (Shanghai Ltd, Shanghai, China) and Peptide 2.0 (USA).

### KIR-Fc binding assay

KIR-Fc binding to cell lines and peptide-loaded cells was assessed by flow cytometry largely as described [19, 21]. KIR2DL1*001-Fc, KIR2DL2*001-Fc and KIR2DL3*001-Fc (R&D, 1844-KR, 3015-KR, 2014-KR) were conjugated to protein-A Alexa Flour 647 (Invitrogen) by overnight incubation at 9:1 (molar) at 4°C, then diluted to 3.6 μg/ml in PBS + 2% fetal calf serum (FCS). 1.25 × 10^5^ cells were placed in 96 flat well plates, resuspended in 25 μl of KIR-Fc (3.6 μg/ml or as indicated) and incubated at 4°C for 1 hr. For peptide loaded cells, KIR-Fc binding was assessed after overnight incubation at 26°C with 100 μM of synthetic peptide. KIR-Fc binding to 221 cells or protein A-Alexa Flour 647 alone were used to baseline values for KIR-Fc binding. Cells were washed with PBS and data acquired by flow cytometry.

### NK cell functional assays

NK cell degranulation assays were carried out largely as described [19]. For Figure 5, Additional file 3G, H and Additional file 6, KIR A/A haplotype donor blood samples from healthy volunteers were recruited and collected under the Imperial College Healthcare NHS Trust Tissue Bank ethical approval (REC number 12/WA/0196). For Figure 6D and Additional file 7D, anonymous donors were recruited from the National Institutes of Health (NIH) Department of Transfusion Medicine under an NIH Institutional Review Board–approved protocol with informed consent. 1x10^6^ isolated PBMCs were incubated overnight at 37°C with 10 ng/ml IL-15 (R&D) and mixed the following day at 5:1 with 221 cell lines or peptide loaded target cells in presence of 1 μl BV421 anti-CD107a mAb for 2hr at 37^o^C (H4A3, BD 562623). PBMCs were stained with mAbs to identify KIR single positive NK cells. NK cells were identified as CD56^+^ (NCAM16.2, BD 562780) and CD3^−^ (SK7, BD 557832). KIR2DL3-SP NK cells were KIR2DL3^+^ (180701, R&D FAB2014F), KIR2DL1^−^ (143211, R&D FAB1844P), KIR3DL1^−^ (DX9, Miltenyi Biotec 130-092-474) and NKG2A^−^ (Z199, Beckman Coulter PNB10246). For Figure 6D and Additional file 7, donors with unknown KIR genotypes were screened for expression for KIR2DL1 and KIR2DS1 with combinations of mAbs (143211 & EB6), as described [50]. Flow cytometry panels were adjusted to allow isolation of KIR2DL1-SP NK cells if the donor carried KIR2DL1 and KIR2DS1. NK cell cytotoxicity assays were carried out with YTS NK cells mixed with 221 target cells at increasing effector target ratios, at 37°C for 2 hr. For experiments with 221–HLA-C+ cell lines, the Europium release assay (Perkin Elmer, USA) was used as described [51]. For assays using peptide loaded cells, propidium iodide incorporation assays were used as described [52].

### Cell lines

721.221 (221) cells and 221 cells expressing HLA-C*04:01 were used (provided by J. Gumperz and P. Parham, Stanford University, USA). The RMA-S cell line expressing HLA-C*08:02 has been described [21]. The 221 and RMA-S cell lines were cultured in RPMI or IMDM (Gibco) supplemented with 10% FCS. YTS cell lines were previously described and cultured in IMDM with 12.5% FCS [51, 53]. HLA-C*05:01 was expressed in 221 cells by retroviral transduction. Briefly, 293T cells stably expressing vesicular stomatitis virus (VSV) gag and pol genes were transiently transfected by calcium phosphate with 6 mg each of VSV-envelope glycoprotein and HLA-C*05:01 cDNAs. The medium was changed at 24 hours and harvested at 48 and 72 hours for infection. On consecutive days, filtered virus-containing supernatant was incubated with untransfected 221 cells and 8 mg/ml polybrene. The plate was centrifuged at 1000 x g, 32°C for 2 hrs. Two days after infection, cells were selected 0.5 mg/ml G418. ICP47 was expressed in 221 cell lines by electroporation as described [53, 54]. Cell lines were cloned by limiting dilution after drug selection.

### Flow Cytometry

KIR-Fc binding, HLA-I stabilization, degranulation, and cytotoxicity assays were acquired on a FACS ARIA II (BD, UK). Instrument identification: BD FACSAria II SORP configuration number 39305 (NIHR BRC FACS and Confocal Imaging Facility, Hammersmith Campus, Imperial College London, UK). Fluidics configuration: 85 mm nozzle, 45 psi, frequency of 45 kHz and a window extension of 2. Optical configuration: 488 nm Laser Line: 780/50, 710/50, 670/14, 610/20, 575/25, 488/10. 633 nm Laser Line: 780/60, 710/50, 670/14. 405 nm Laser Line: 610/20, 525/50, 450/40. Data were exported as FCS files and analyzed using FlowJo software (Treestar, version 10). Compensation for multi-color experiments were set using single mAb stained beads and Cytometer Setup and Tracking beads were run daily.

### Human donors

All studies were conducted according to the principles expressed in the Declaration of Helsinki. Healthy volunteers were recruited with informed consent with ethical approval either through the Imperial College Human Tissue Bank, London (REC number 12/WA/0196) or from the National Institutes of Health (NIH) Department of Transfusion Medicine under an NIH Institutional Review Board–approved protocol with informed consent.

### Plasmids

An HLA-C*05:01 cDNA [55] was supplied by Pierre Coulie, as plasmid pcDNA3.1–HLA-C*05:01 (de Duve Institute, Université Catholique de Louvain, UCL, Brussels, Belgium). Ten His residues followed by a stop codon were added to the C-terminus of HLA-C*05:01 by PCR of the HLA-C*05:01 cDNA. HLA-C*05:01-10xHis was excised with BamHI and EcoRI and cloned into pBABE-puro-CMV^+^ by blunt-end ligation via the SalI site. pzLRS-ICP47-IRES-GFP [56] was supplied by Emmanuel Wiertz (Center of Infectious Diseases and Department of Medical Microbiology, Leiden University Medical Center, Leiden, The Netherlands). HLA-C*08:02 cDNA was generated by two single base changes in codons 77 and 80 of HLA-C*05:01 cDNA (Fig. 1A) by Quickchange PCR of HLA-C*05:01 using the primers

Fwd: 5’-CAGACTGACCGAGTGAGCCTGCGGAACCTGCGCGGCTACTAC-3’ and Rev: 5’- GTAGTAGCCGCGCAGGTTCCGCAGGCTCACTCGGTCAGTCTG-3’.

### Statistical analysis

All statistical analysis was carried out in GraphPad PRISM (version 5.0). Multiple groups were compared by Kruskal-Wallis test with Dunn’s multiple comparisons test.

## Abbreviations

CD3: cluster of differentiation 3
CHAPS: 3-[(3-Cholamidopropyl) dimethyl ammonio]-1-propanesulfonate
CMV: Cytomegalovirus
EBV: Epstein–Barr virus
EDTA: Ethylenediaminetetraacetic acid
FACS: Fluorescence activated cell sorting
GFP: Green fluorescent protein
HCV: Herpes simplex virus
HIV: Human immunodeficiency virus
HLA: Human leukocyte antigen
KIR: Killer-cell immunoglobulin-like receptor
MFI: Median fluorescence intensity
NK: cell Natural killer cell
PBMC: Peripheral blood mononuclear cell
PBS: Phosphate-buffered saline
SEM: Standard error of the mean
SP: single positive
TAP: Transporter associated with antigen processing
VSV: Vesicular stomatitis virus

## Declarations

### Ethics approval and consent to participate

Blood samples from healthy volunteers for this study were recruited and collected under the Imperial College Healthcare NHS (National Health Service) Trust Tissue Bank ethical approval (REC number 12/WA/0196), and from the NIH (National Institutes of Health) Department of Transfusion Medicine, under an NIH Institutional Review Board-approved protocol, with informed consent.

### Authors’ contributions

M.J.W.S designed the study, developed and performed experiments, interpreted data and prepared the manuscript. S.R, M.E.P, and A.K performed experiments. J.M.S recruited healthy controls for the study. S.A.M, J.S and D.F.H carried out mass spectrometry analysis. R.J.B, D.M.A and E.O.L conceived and designed the study, interpreted the data, and prepared the manuscript. R.J.B and E.O.L. supervised the research and contributed equally to the study. All authors discussed the results and commented on the manuscript.

## Acknowledgements

We thank M. Opanowicz, J. Manji, L. Black, J. Roberts, and the Imperial College London Hammersmith Campus NIHR BRC Flow Cytometry and Confocal Imaging Facility for assistance with flow cytometry. We thank P. Coulie and E. Wiertz for plasmids, and J. Gumperz and P. Parham for cell lines.

## Additional Files

**Additional file 1.**
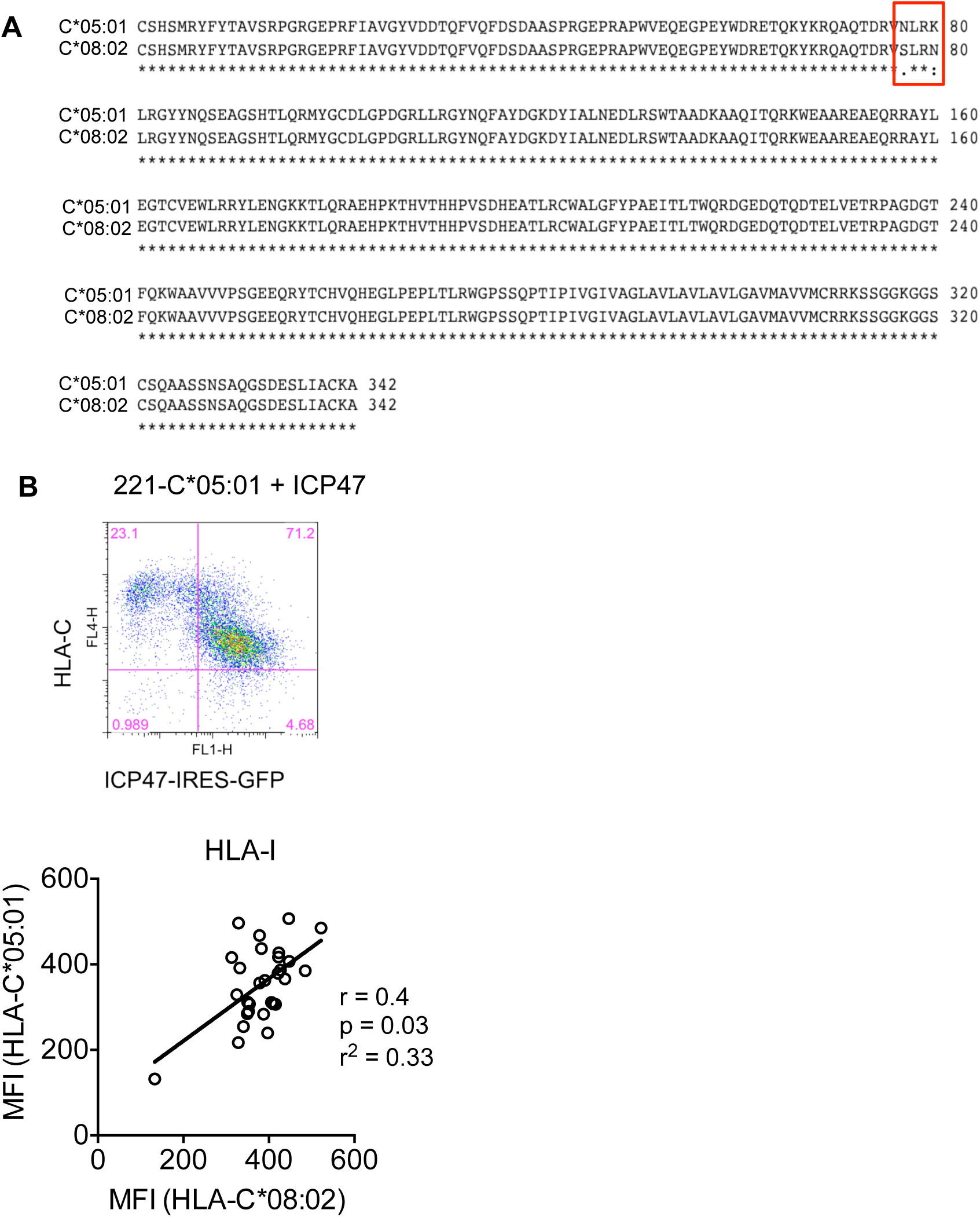
Sequence alignment of HLA-C*05:01 and HLA-C*08:02, generation of TAP-deficient cell line expressing HLA-C*05:01 and correlation of HLA-I stabilization. **(A)** Amino acid alignment of HLA-C*05:01 and HLA-C*08:02. Box indicates positions 77-80. **(B)** Expression of ICP47 (via IRES-GFP) and HLA-C in 221–C*05:01 cells transfected with pzLRS– ICP47-IRES-GFP. After drug selection, 221–C*05:01–ICP47 cell line was cloned by limiting dilution. **(C)** Correlation of HLA-I stabilization. HLA-C*05:01 against HLA-C*08:02 (Spearman).

**Additional file 2.**
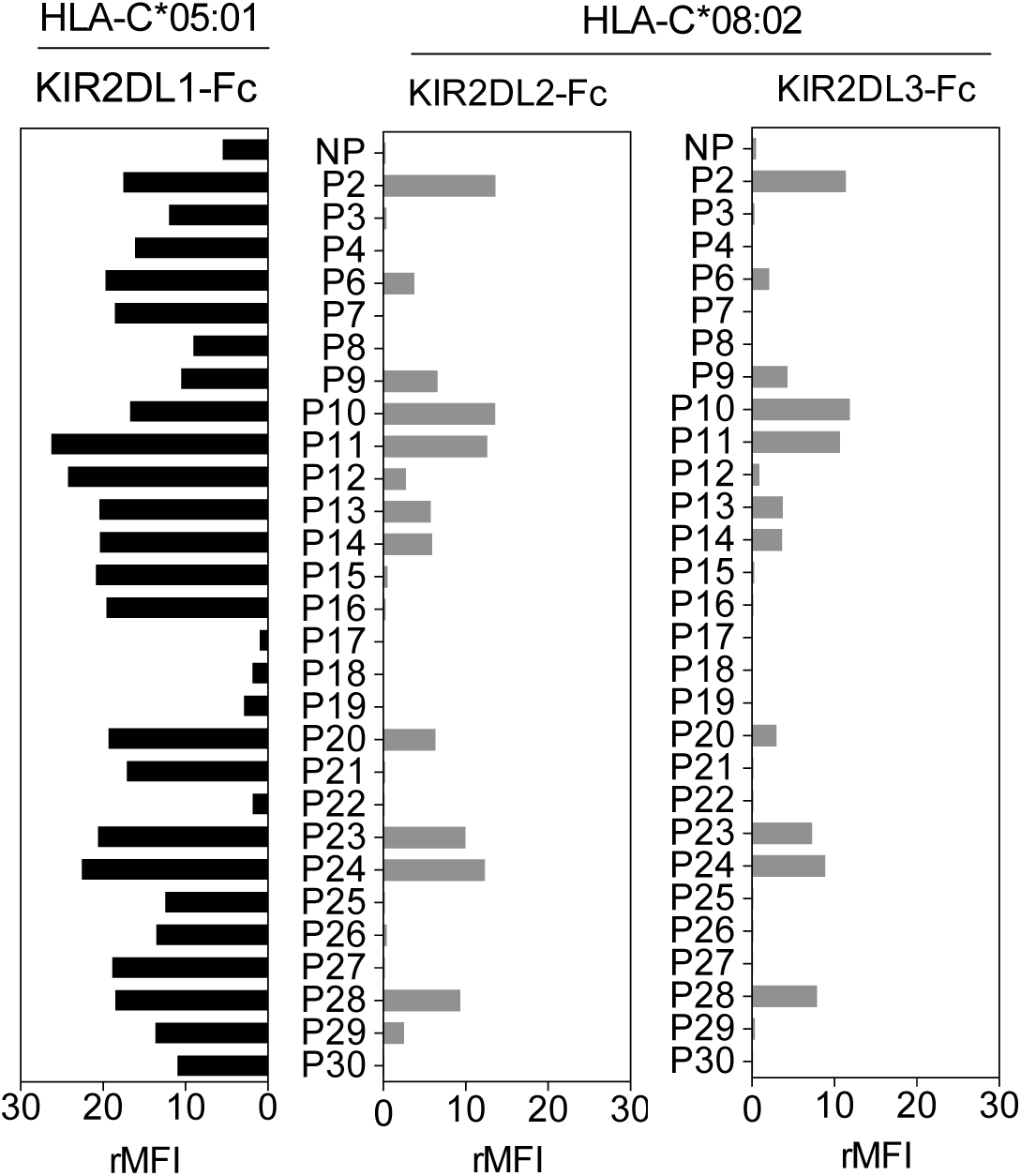
KIR2DL2 and KIR2DL3 binding to HLA-C*08:02 is more peptide selective than KIR2DL1 binding to HLA-C*05:01. Data from Figure 2B normalized to HLA-I (KIF-Fc MFI/HLA-I MFI) for each peptide.

**Additional file 3.**
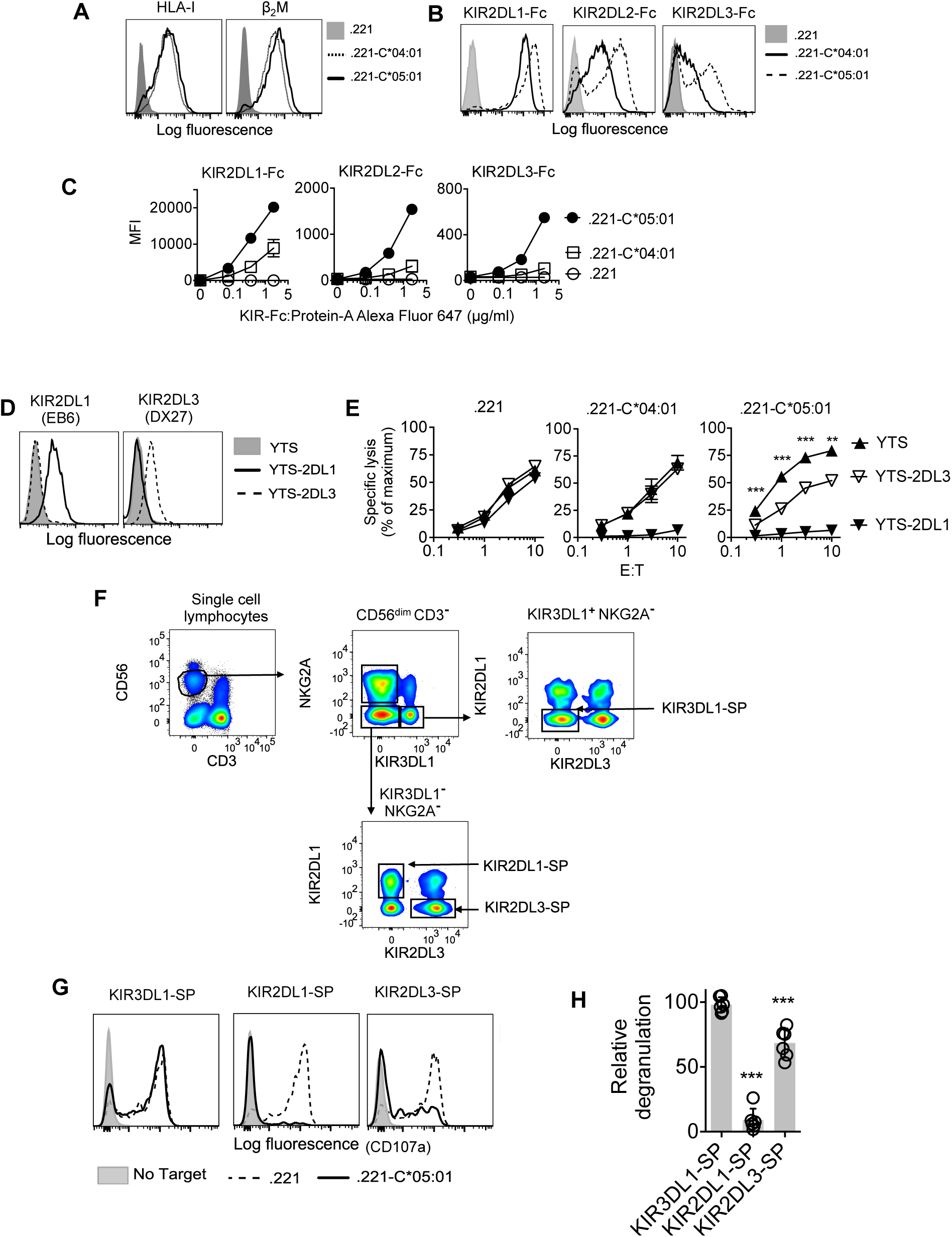
HLA-C*05:01 is a functional ligand for KIR2DL1, KIR2DL2 and KIR2DL3. **(A)** Expression of HLA-I and β2 microglobulin on 221, 221–C*04:01 and 221–C*05:01 cells. **(B** and C) KIR2DL1-Fc, KIR2DL2-Fc and KIR2DL3-Fc binding to 221, 221–C*04:01 and 221–C*05:01 cells as determined by flow cytometry. **(C)** Mean MFI and SEM are shown from three independent experiments. **(D)** Expression of KIR2DL1 and KIR2DL3 on YTS, YTS-2DL1 and YTS-2DL3 NK cells. **(E)** Specific lysis of 221, 221–C*04:01 and 221–C*05:01 cells by YTS, YTS-2DL1 and YTS-2DL3 NK cells at different effector-target ratios (E:T). Mean and SEM of three independent experiments are shown. **=p<0.01, ***=p<0.001 by students t test. **(F)** Flow cytometry gating strategy to identify KIR single positive (KIR-SP) NK cells. CD56dim CD3^−^ lymphocytes were identified from PBMCs. KIR2DL3-SP NK cells were identified as KIR3DL1^−^ and NKG2A^−^, KIR2DL1^−^ NK cells from A/A KIR haplotype donors. **(G)** Flow cytometry histograms showing degranulation (CD107a at the cell surface) of KIR3DL1 single positive (SP), KIR2DL1-SP and KIR2DL3-SP NK cells in response to 221, 221–C*05:01 or no target cells. **(H)** Degranulation of KIR-SP NK cells in response to 221 and 221–C*05:01 cells for N=5 A/A KIR haplotype donors. Degranulation is shown relative to 221. \*\*\**P*<0.001 by students t test.

**Additional file 4.**
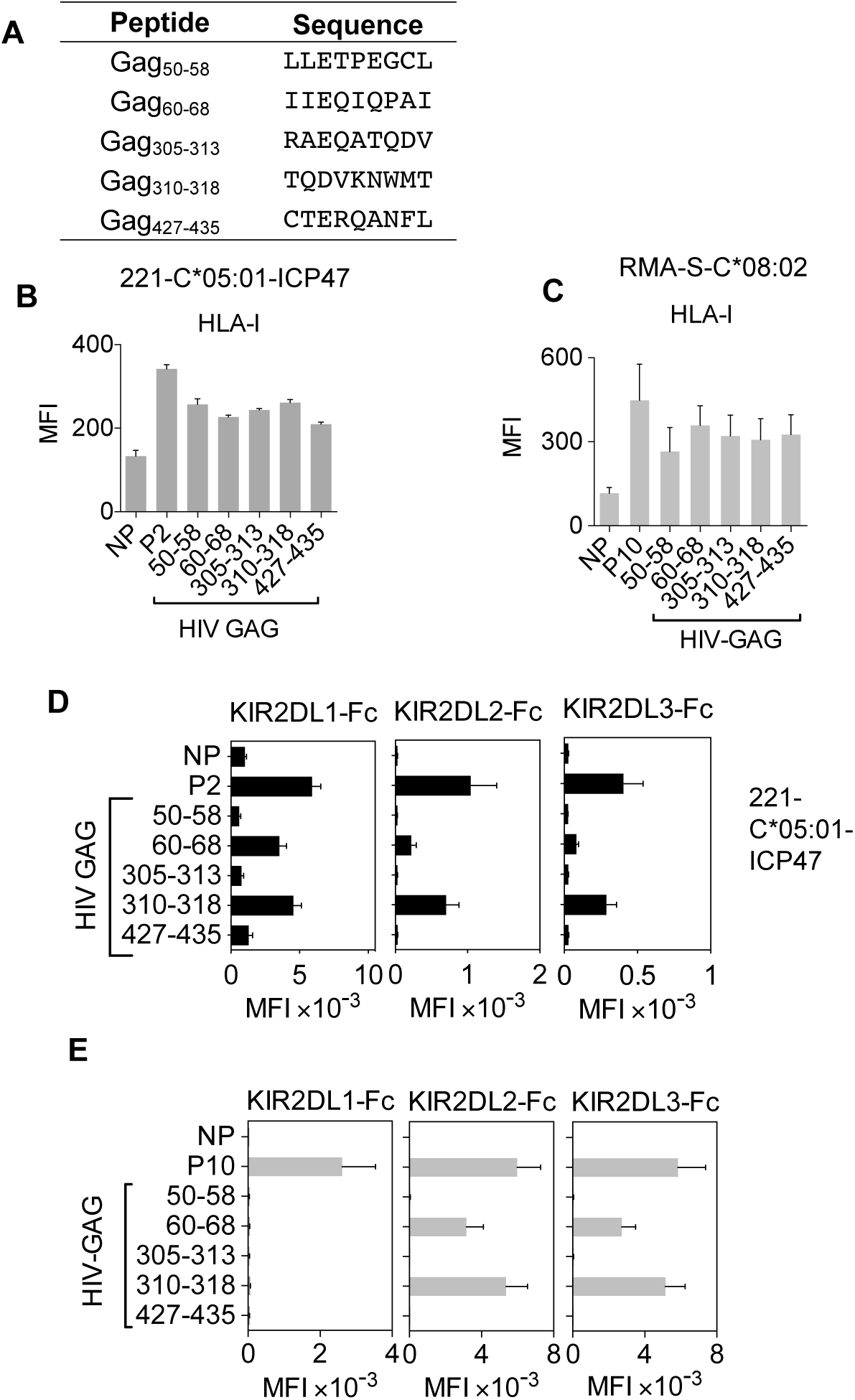
HLA-C*05:01 HIV Gag peptides modulate canonical and cross-reactive binding of KIR2DL2 and KIR2DL3 to HLA-C*05:01(C2). **(A)** HLA-C*05:01 HIV Gag 9mer peptide sequences. **(B)** Expression of HLA-I on 221–C*05:01–ICP47 cells loaded with no peptide (NP), P2 and HIV Gag peptides 50-58, 60-68, 305-313, 310-318 and 427-435. Mean MFI and SEM of 3 independent experiments are shown. **(C)** Expression of HLA-I on RMA-S-C*08:02 cells loaded with no peptide (NP), P2 and HIV Gag peptides 50-58, 60-68, 305-313, 310-318 and 427-435. Mean MFI and SEM of 3 independent experiments are shown. **(D)** KIR2DL1-FC, KIR2DL2-Fc and KIR2DL3-Fc binding to 221–C*05:01–ICP47 cells loaded with NP, P2 or HIV Gag peptides, 50-58, 60-68, 305-313, 310-318 and 427-435. Mean MFI and SEM of 3 independent experiments are shown. **(E)** KIR2DL1-Fc, KIR2DL2-Fc and KIR2DL3-Fc binding to RMA-S-C*08:02 cells loaded with NP, P2 or HIV Gag peptides, 50-58, 60-68, 305-313, 310-318 and 427-435. Mean MFI and SEM of 3 independent experiments are shown.

**Additional file 5.**
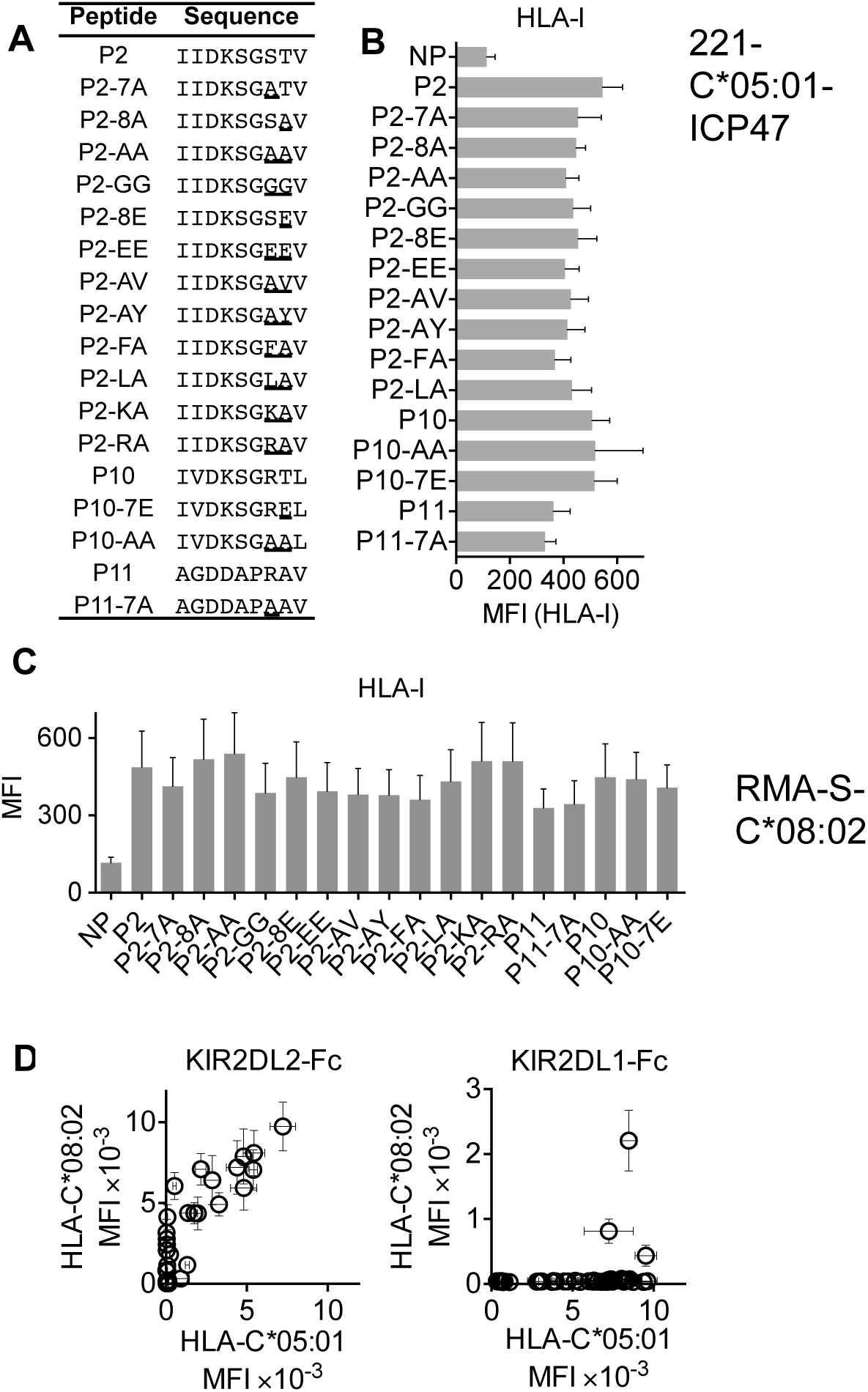
Stabilization of HLA-I on TAP-deficient cells by peptides and crossreactive binding of KIR2DL2 to peptide-loaded HLA-C*05:01 and KIR2DL1 to peptide-loaded HLA-C*08:02. **(A)** Sequences of peptides P2, P10, P11 and peptides with amino acid substitutions at positions 7 and 8. **(B)** HLA-I expression on 221–C*05:01–ICP47 cells after overnight culture in the presence of HLA-C*05:01 peptides P2, P10, P11 and amino acid substituted peptides compared to cells with no peptide (NP). Mean MFI and SEM of three independent experiments are shown. **(C)** HLA-I expression on RMA-S-C*08:02 cells after overnight culture in the presence of HLA-C*05:01 peptides P2, P10, P11 and amino acid substituted peptides compared to cells with no peptide (NP). Mean MFI and SEM of three independent experiments are shown. **(D)** KIR2DL2-Fc *(left)* and KIR2DL1-Fc *(right)* binding to HLA-C*05:01 is correlated with binding to HLA-C*08:02 in the presence of the same peptides. All peptides with amino acid substitutions are shown.

**Additional file 6.**
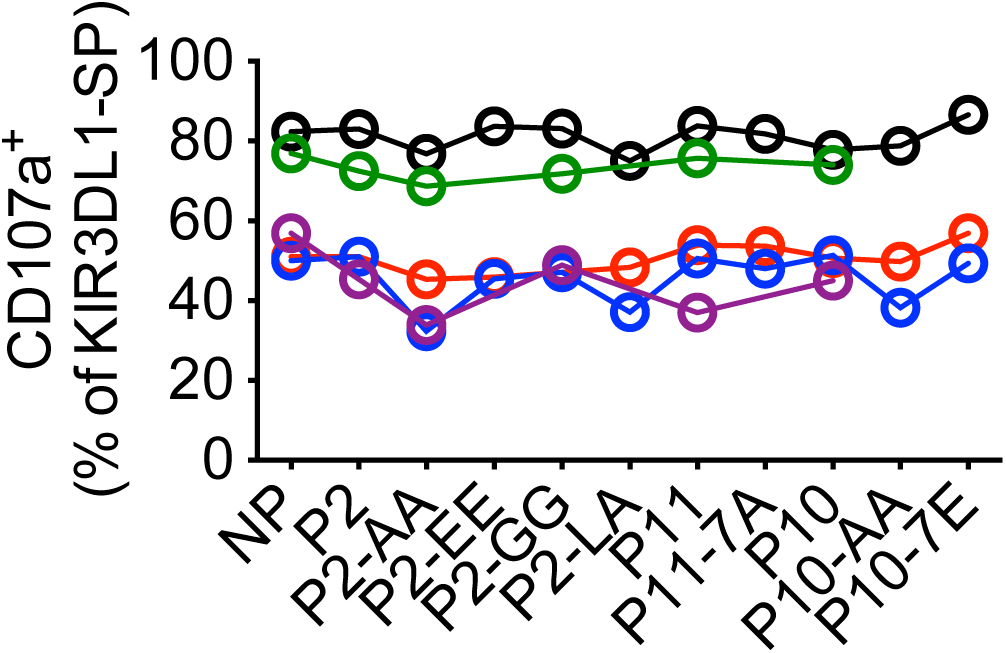
Degranulation of KIR3DL1-SP NK cells in response to peptide-loaded TAP-deficient HLA-C*05:01 cells. KIR3DL1-SP NK cell CD107a expression (% positive) in response to 221–C*05:01–ICP47 cells loaded with no peptide (NP), P2, P2-AA, P2-EE, P2-LA, P2-GG, P11, P11-7A, P10, P10-AA and P10-7E. Individual donors are represented by different colours.

**Additional file 7.**
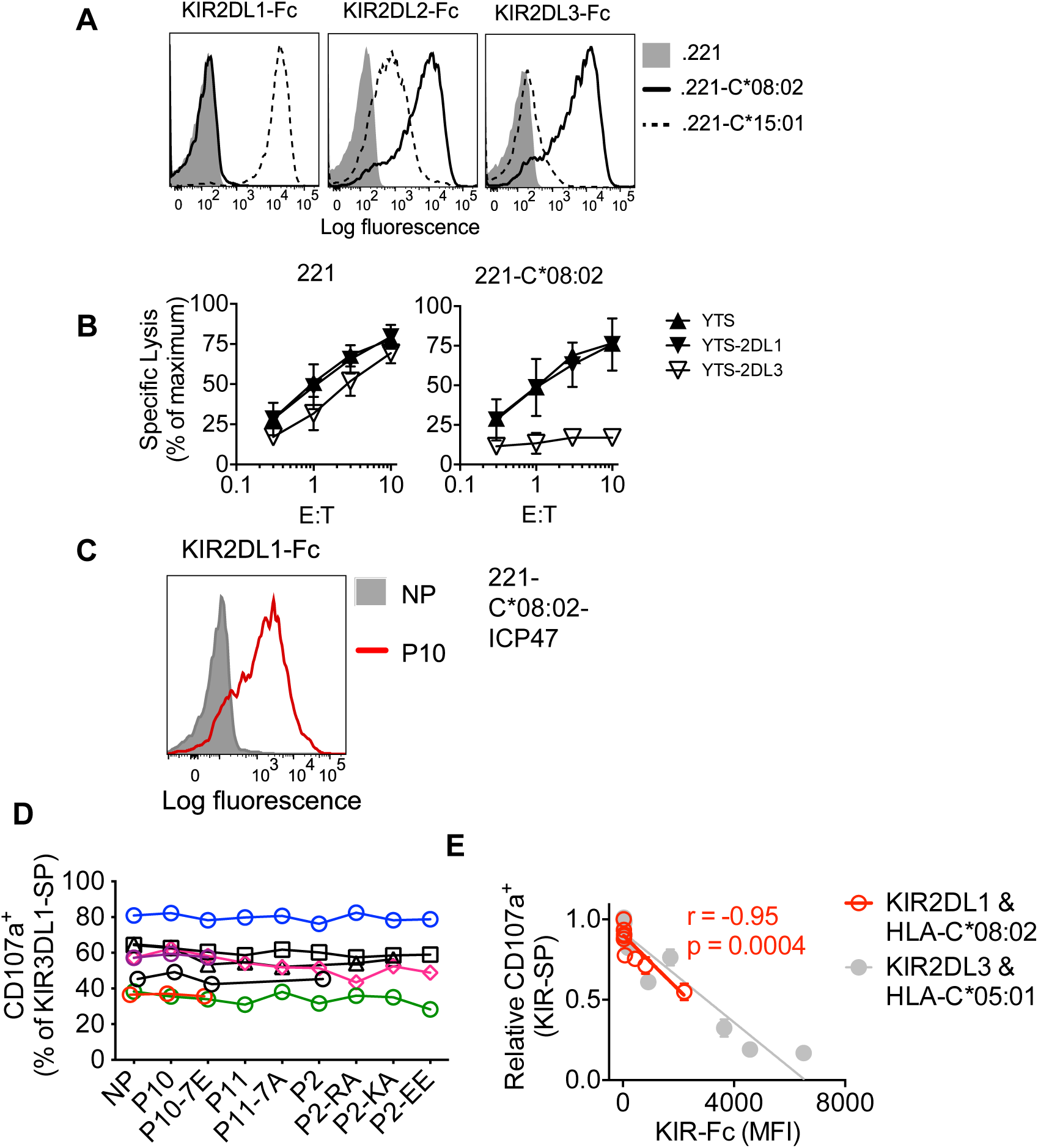
KIR-Fc binding and functional responses of KIR^+^ NK cells to 221-C*08:02 and 221-C*08:02-ICP47 cells. (**A**) KIR2DL1-Fc, KIR2DL2-Fc and KIR2DL3-Fc binding to 221, 221–C*08:02 and 221–C*15:01. (**B**) Specific lysis of 221 and 221–C*08:02 cells by YTS, YTS-2DL1 and YTS-2DL3 NK cells. (**C**) KIR2DL1-Fc binding to 221–C*08:02–ICP47 cells loaded with no peptide (NP) or P10. (**D**) KIR3DL1-SP NK cell CD107a expression (% positive) in response to 221–C*08:02–ICP47 cells loaded with no peptide (NP), P2, P2-AA, P2-EE, P2-LA, P2-GG, P11, P11-7A, P10, P10-AA and P10-7E. Individual donors are represented by different colours. (**E**) Overlay of cross-reactive KIR-SP NK cell responses and cross-reactive KIR-Fc binding to HLA-C*05:01 and HLA-C*08:02.

